# Open access-enabled evaluation of epigenetic age acceleration in colorectal cancer and development of a classifier with diagnostic potential

**DOI:** 10.1101/2023.08.29.555284

**Authors:** Tyas Arum Widayati, Jadesada Schneider, Kseniia Panteleeva, Elizabeth Chernysheva, Natalie Hrbkova, Stephan Beck, Vitaly Voloshin, Olga Chervova

**Affiliations:** Medical Genomics Lab, Cancer Institute, University College London, London, United Kingdom; Department of Genetics, Evolution and Environment, University College London, London, United Kingdom; Department of Pathology and Biomedical Science, University of Otago, Christchurch, New Zealand; School of Biological and Behavioural Sciences, Queen Mary University of London, London, United Kingdom

**Author notes:** Correspondence: Tyas Arum Widayati, Olga Chervova.

**Keywords:** epigenetic age, colorectal cancer, CRC, epigenetic clock, epigenetic age acceleration, colon tissue methylation

## Abstract

Aberrant DNA methylation (DNAm) is known to be associated with the aetiology of cancer, including colorectal cancer (CRC). In the past, the availability of open access data has been the main driver of innovative method development and research training. However, this is increasingly being eroded by the move to controlled access, particularly of medical data, including cancer DNAm data. To rejuvenate this valuable tradition, we leveraged DNAm data from 1,845 samples (535 CRC tumours, 522 normal colon tissues adjacent to tumours, 72 colorectal adenomas, and 716 normal colon tissues from healthy individuals) from 14 open access studies deposited in NCBI GEO and ArrayExpress. We calculated each sample’s epigenetic age (EA) using eleven epigenetic clock models and derived the corresponding epigenetic age acceleration (EAA). For EA, we observed that most first- and second-generation epigenetic clocks reflect the chronological age in normal tissues adjacent to tumours and healthy individuals (e.g. Horvath (*r* = 0.77 and 0.79), Zhang EN (*r* = 0.70 and 0.73)) unlike the epigenetic mitotic clocks (EpiTOC, HypoClock, MiAge) (*r <* 0.3). For EAA, we used PhenoAge, Wu, and the above mitotic clocks and found them to have distinct distributions in different tissue types, particularly between normal colon tissues adjacent to tumours and cancerous tumours, as well as between normal colon tissues adjacent to tumours and normal colon tissue from healthy individuals. Finally, we harnessed these associations to develop a classifier using elastic net regression (with lasso and ridge regularisations) that predicts CRC diagnosis based on a patient’s sex and EAAs calculated from histologically normal controls (i.e. normal colon tissues adjacent to tumours and normal colon tissue from healthy individuals). The classifier demonstrated good diagnostic potential with ROC-AUC=0.886, which suggests that an EAA-based classifier trained on relevant data could become a tool to support diagnostic/prognostic decisions in CRC for clinical professionals. Our study also reemphasises the importance of open access clinical data for method development and training of young scientists. Obtaining the required approvals for controlled access data would not have been possible in the timeframe of this study.

## 1 INTRODUCTION

Colorectal cancer (CRC) is the third most common cancer in the world, with around 1.93 million new cases worldwide in 2020 (Sung et al. (2021)). One of the main risk factors of CRC is ageing (Dekker et al. (2019)). Here, ageing is not solely referred to as an increase in chronological age (CA), but is also viewed as a gradual decline in biological function (biological ageing) (Gems (2015)). One of the hallmarks of ageing is epigenetic alteration, which includes changes in DNA methylation (DNAm) patterns, abnormal histone modifications, and irregular chromatin remodelling (López-Otín et al. (2013)). Epigenetic alteration is one of the hallmarks of cancer, including CRC (Dekker et al. (2019); Hanahan (2022)). CRC arises due to the accumulation of genetic and epigenetic alterations in the colon mucosa. Abnormal changes in DNAm patterns are a common form of epigenetic change in CRC. They contribute to the initiation of abnormal stem cell growth of the intestine, this is often followed by the appearance of adenomas and, later, progression to carcinoma (Dekker et al. (2019); Schmitt and Greten (2021)). Interestingly, DNAm alteration was not only observed in cancerous tissues but also in normal colon tissue, indicating the early occurrence of DNAm changes in CRC tumour development or the field effect of cancerisation (Luo et al. (2014); Joo et al. (2021); Sanz-Pamplona et al. (2014)).

There are several methods developed for CRC diagnosis, with colonoscopy being considered the gold standard (Dekker et al. (2019)). Yet, other potential prognostic and diagnostic markers, including DNAm-based biomarkers, have been studied in order to provide robust results (Okugawa et al. (2015); Mueller and Gyoőrffy (2022)). DNAm pattern abnormalities in cancer, including in CRC, occur due to hyper- and/or hypo-methylation of some genomic regions (Nishiyama and Nakanishi (2021)). Some CRC cases are also associated with a unique CpG island methylator phenotype (CIMP), which is characterised by the strong hypermethylation in certain promoter regions across the genome (Schmitt and Greten (2021)).

In the past decade, epigenetic age predictors (”epigenetic clocks”) have been developed to estimate chronological and biological age based on DNAm levels in specific age-associated CpG sites (Table 1). The first-generation epigenetic clocks, namely Horvath and Hannum clocks, were mainly utilised to predict chronological age (Horvath (2013); Hannum et al. (2013)). Second-generation clocks were then developed to not only estimate the chronological age but also to capture physiological conditions by incorporating some clinical measures (e.g. blood biomarkers) or by including specific CpG sites in their models (Levine et al. (2018); Horvath et al. (2018); Wu et al. (2019); Zhang et al. (2019)). Later, some cancer-specific epigenetic clock models were constructed by combining molecular mitotic clocks and cancer DNAm pattern alteration hypotheses (Yang et al. (2016); Youn and Wang (2018); Teschendorff (2020)).

**Table 1.** Summary of the epigenetic clocks. Abbreviations: DNAm - DNA methylation, CpG - cytosine phosphate guanine

| Category | Clocks (reference) | Description |
| --- | --- | --- |
| First-generation clocks | Horvath (Horvath (2013)) | Developed on DNAm of various tissue samples. Used penalised regression model to regress CA onto 353 CpG sites (which are previously selected by elastic net (EN) regression model). |
|  | Hannum (Hannum et al. (2013)) | Developed by regressing CA onto blood DNAm data using EN regression model, which resulted in selected 71 CpG sites as the accurate CA predictor. |
| Second-generation clocks | PhenoAge (Levine et al. (2018)) | Developed through two-step process: determination of "phenotypic age" metric and regression of blood DNAm data onto phenotypic age, resulting in selected 513 CpG sites to estimate final phenotypic age. |
|  | Skin and Blood (Horvath et al. (2018)) | This clock uses 391 CpGs to estimate epigenetic age. These CpGs were obtained from EN regression of CA onto blood DNAm, saliva, fibroblasts, keratinocytes, buccal cells, and endothelial cells. |
|  | Pediatric-Buccal-Epigenetic (PedBE) (McEwen et al. (2020)) | This clock uses 94 CpG sites to predict epigenetic age. Elastic net regression on pediatric buccal DNAm data was used to select these CpG sites. |
|  | Wu (Wu et al. (2019)) | Trained on paediatric blood DNAm from 11 datasets. Elastic net approach used in this model resulted in selected 111 CpG sites to estimate child-specific biological age. |
|  | Zhang BLUP (Zhang et al. (2019)) | Trained on blood and saliva DNAm. Uses 319,607 CpG probes (obtained using Best Linear Unbiased Prediction (BLUP) approach) to estimate epigenetic age. |
|  | Zhang EN(Zhang et al. (2019)) | Trained on blood and saliva DNAm. Uses 514 CpG sites (selected using EN regression) to estimate epigenetic age. |
| Epigenetic mitotic clocks | EpiTOC (Yang et al. (2016)) | This clock uses average DNAm level of 385 CpGs from PCGT promoters that are generally unmethylated in 11 foetal tissue types to predict mitotic age. |
|  | HypoClock (Teschendorff (2020)) | This clock uses average DNAm level of 678 solo-WCGW sites. |
|  | MIage (Youn and Wang (2018)) | Trained on 4,020 cancer and adjacent normal tissue DNAm from 8 TCGA cancer data, and tested on 5 other TCGA cancer data. Used the panel of selected 268 hypermethylated CpGs to estimate mitotic age. |

Deviation of the predicted epigenetic age (EA) from the chronological age (CA), known as epigenetic age acceleration (EAA), has been studied with respect to its association with age-related phenotypic changes and health outcomes, including cancer (Horvath (2013); Oblak et al. (2021)). Since DNAm alteration is associated with cancer incidence, epigenetic age scores have been studied to find suitable DNAm markers for cancer, including CRC. Previous studies have assessed the relationship between CRC and EAA (Durso et al. (2017); Zheng et al. (2019); Devall et al. (2021); Nwanaji-Enwerem et al. (2021); Matas et al. (2022)). However, our understanding of whether epigenetic ageing measures (EA and/or EAA) differ between histologically normal colon tissues in individuals with and without CRC is limited to two publications (Joo et al. (2021); Wang et al. (2020)). These studies identified a significant difference in epigenetic age acceleration between normal colon tissue from patients with and without CRC. However, although both studies assessed the same clocks (i.e., Horvath, Hannum, PhenoAge, EpiTOC), they obtained different results. Joo et al. (2021) found a significant difference in EpiTOC age acceleration while Wang et al. (2020) observed it in EAA from the PhenoAge clock. The differences in datasets, sample groupings, and number of samples in each study may be a plausible explanation for this. Hence, to identify the most suitable clock for reflecting DNAm changes in CRC, further study regarding the associations between epigenetic clock measures and CRC, particularly in normal colon tissue, is needed.

This study was designed to be suitable for a Masters’s student project (i.e., it had to be completed within six months). Although the vast majority of DNAm data, including for CRC, are deposited in public databases such as EGA and dbGaP, they are classified as controlled access data which requires a data access agreement to be completed and to be approved by a data access committee before the data can be shared. This process can take months or even years (Powell (2021)) and is further complicated by diverse and, in some cases, even inappropriate data access agreements (Saulnier et al. (2019)). For these reasons, only data that are available under open access were considered for inclusion in this study. Despite being rare, open access data are of equal quality and have a long and successful track record as drivers of innovation and training (Greenbaum et al. (2011)). The resulting limitations and advantages of using exclusively open access data are discussed further in Section 4.3.

We obtained 14 open access datasets (summarised in Table S1) with the aim of evaluating the associations between CRC diagnosis and epigenetic ageing measures (EAs and EAAs) derived from eleven epigenetic clocks. In particular, we aimed to: (1) evaluate the associations between chronological age and estimated EAs for each tissue type; (2) identify the EAAs that can capture the difference between CRC tumours, normal colon tissues adjacent to tumours, colorectal adenomas, and normal colon tissues from healthy individuals; (3) determine the EAAs that can distinguish between histologically normal colon tissues from individuals with different CRC diagnoses; and (4) develop an EAA-based classifier that demonstrates good potential for use in distinguishing between normal colon tissues from healthy individuals and normal colon tissues adjacent to tumours, thus aiding CRC diagnosis. Graphical overview of the study design is presented on Figure 1, the methodology is summarised in Figure S1.

**Figure 1.**
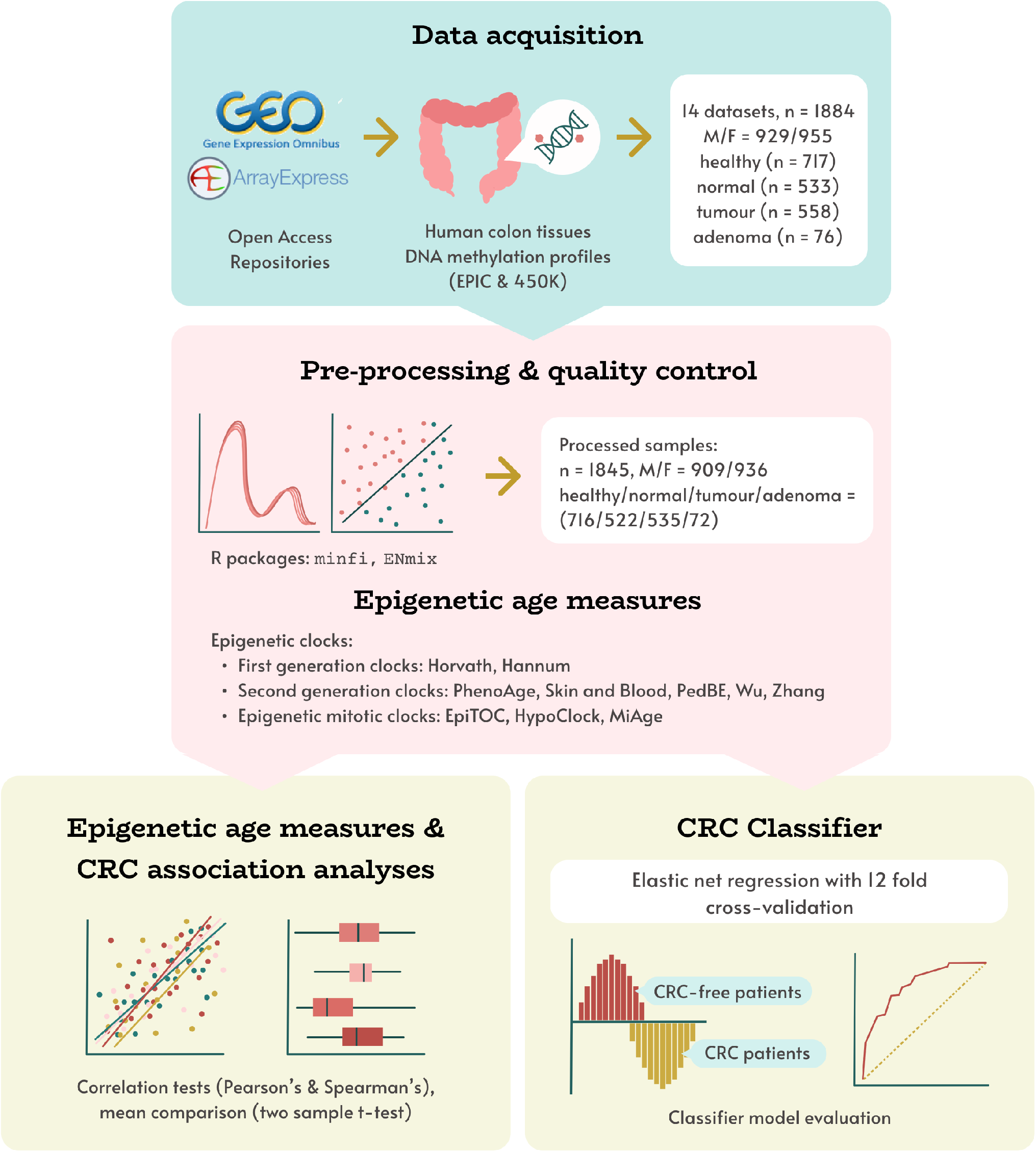
Study design overview. Human colon DNAm datasets, obtained from open access repositories was pre-processed, and corresponding epigenetic age measures were calculated using 11 DNAm clocks. These measures were used in evaluating associations between epigenetic age and age acceleration with tissue type (healthy, normal, adenoma, tumour), and developing a novel CRC status classifier model.

## 2 METHODS

### 2.1 Association analysis

#### 2.1.1 Data acquisition and pre-processing

The data for this study were downloaded from two public repositories: NCBI GEO (National Center for Biotechnology Information Gene Expression Omnibus) and EMBL-EBI (European Molecular Biology Laboratory European Bioinformatics Institute) ArrayExpress (Barrett et al. (2012); Sarkans et al. (2021)). The list of datasets used in this study is given in Table S1. In particular, we searched for human colon tissue DNA methylation (DNAm) profiles generated using Illumina methylation platforms (Infinium HumanMethylation450 and MethylationEPIC arrays), with available chronological age, colorectal cancer (CRC) patient status, and specimen pathology (tumour, adenoma or normal tissue) (Bibikova et al. (2011); Pidsley et al. (2016)). Dataset GSE132804, which includes DNAm profiles produced using both 450K and EPIC platforms, was treated as two separate datasets with respect to the technology used.

Where possible, the data were processed from raw .idat files for each dataset separately following previously described methods (Chervova et al. (2019)). In brief, samples with more than 1% of low-quality probes (detection *p >* 0.01, bead count *<* 3), or in disagreement between reported and inferred sex, were excluded, together with samples identified as outliers by built-in quality control checks of `minfi` and `ENmix` R packages (Aryee et al. (2014); Xu et al. (2021); R Core Team (2009)). Missing and low-quality CpG probes (across more than 1% of samples) were filtered out. Data were normalised using the ssNoob method implemented in the `minfi` package (Fortin et al. (2017)). For some datasets without raw data and/or necessary technical information, we used published pre-processed data and performed quality control checks by assessing their methylation values data (distribution plots, reported and inferred sex matches).

#### 2.1.2 Sample notations and variables description

All samples in our data contain information regarding chronological age, sex, and tissue types. We categorised samples into four different tissue types:

- **healthy:** samples from normal colon tissues of individuals without CRC (i.e. no concurrent CRC was observed at the time of sample collection); normal colon tissues from individuals with concurrent colon adenoma were included in this category,
- **normal:** samples from normal colon tissues adjacent to the tumours of CRC patients,
- **tumour:** samples from cancerous tumours obtained from CRC patients,
- **adenoma:** samples from adenoma tissues of patients with observed colorectal adenoma (mostly sessile serrated adenomas).

For association analysis, we used two different datasets: (a) dataset with healthy, normal, tumour, and adenoma samples (Dataset 1) and (b) dataset with only healthy and normal samples (Dataset 2). A summary of the available cohort characteristics is given in Table 2). Details about the sample collection site (i.e. left or right colon) are available for only half of the dataset. Some samples also have information regarding the detailed location. We classified samples from descending colon, rectosigmoid junction, rectum, sigmoid, and splenic flexure as samples from the left colon, while ascending colon, caecum, hepatic flexure, and transverse colon are from the right colon (Lin et al. (2016). Other information such as race/ethnicity, cancer stage, mutation, and CpG island methylator phenotype (CIMP) status is limited to a small number of samples, hence we excluded these variables from the analysis.

**Table 2.** Summary of cohort characteristics.

|  | Dataset 1 |  |  |  |  | Dataset 2 |  |  |
| --- | --- | --- | --- | --- | --- | --- | --- | --- |
|  | All | Healthy | Normal | Tumour | Adenoma | All | Healthy | Normal |
| No. of samples | 1845 | 716 | 522 | 535 | 72 | 1220 | 715 | 505 |
| Age<br>(median (range)<br>in years) | 63<br>(25.1 - 93.6) | 59<br>(31 - 88) | 64<br>(25.1 - 93) | 66<br>(27 - 93.6) | 75<br>(50 - 90) | 60<br>(25.1 - 93) | 59<br>(31 - 88) | 64<br>(25.1 - 93) |
| Gender |  |  |  |  |  |  |  |  |
| Female | 936 | 453 | 206 | 229 | 48 | 650 | 453 | 197 |
| Male | 909 | 263 | 316 | 306 | 24 | 570 | 262 | 308 |
| Site |  |  |  |  |  |  |  |  |
| Left | 637 | 426 | 140 | 71 | 0 | 561 | 426 | 135 |
| Right | 307 | 218 | 46 | 43 | 0 | 263 | 217 | 46 |
| NA | 901 | 72 | 336 | 421 | 72 | 396 | 72 | 324 |

#### 2.1.3 Epigenetic age calculation

We classified the epigenetic clocks into three categories: first-generation, second-generation, and epigenetic mitotic clocks. First- and second-generation epigenetic age (EA) were calculated for each sample using R `methylClock` library (Pelegí-Sisó et al. (2021)), while epigenetic mitotic clocks were run using the scripts provided by their authors (Yang et al. (2016); Youn and Wang (2018); Teschendorff (2020)). Estimated age and mitotic age scores were used to calculate epigenetic age acceleration (EAA) which is described in the next section. Further details about the epigenetic clocks and EAAs are provided in Table 1.

#### 2.1.4 EAA calculation and statistical analysis

We performed the analysis of outliers separately for Dataset 1 and Dataset 2 by using the differences between epigenetic and chronological age values, which we call epigenetic age acceleration differences (EAAd). This metric was only calculated for the first- and second-generation clocks, and not for the mitotic clocks. A sample was labelled an outlier if its EAAd value was more than three standard deviations away from the mean EAAd across the whole dataset (i.e., outside the interval mean ± 3·SD). We removed all samples which were outliers in at least two clocks. In total, 142 and 38 samples were removed as outliers from Dataset 1 and Dataset 2, respectively.

All analyses in this study were conducted in R v. 4.2.2 (R Core Team (2009)). To evaluate the associations between EAA and CRC, we calculated EAAs from each epigenetic clock using the following steps (EAA for Dataset 1 and Dataset 2 were calculated separately using the same steps):

- **Step 1a:** We regressed epigenetic age onto the chronological age and sex of healthy samples using the linear model (1).

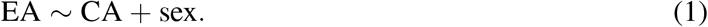

Healthy samples were chosen to ensure the uniform EAA calculation for all epigenetic age scores, including those for mitotic clocks.
- **Step 2a:** Using the linear regression coefficients obtained in Step 1a in model (1), we calculated EAAs as the model residuals.
- **Step 3a:** Based on the mixed-effect model (2), we adjusted EAAs obtained in Step 2a for the dataset and patient IDs using formula (2). This adjustment was made to ensure data independence because in some datasets there is more than one sample per patient, and without this adjustment, they would violate the independence assumption of most statistical tests. Adjustment for dataset ID is to alleviate any batch effect.

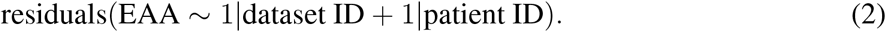

It is worth noting that traditionally EAAs for the first- and second-generation epigenetic clocks are calculated either as differences between EA and CA or as the residuals from linear regression of EA onto chronological age using the whole dataset (Horvath (2013); McEwen et al. (2020)). This works well when the output of the epigenetic clock is predicted age, which correlates well with chronological age. Epigenetic mitotic clocks predict the number of cell divisions (as a proxy to the quality of maintenance of ageing cells). The residuals from fitting mitotic predicted “age” to CA are much less interpretable, as they cannot be easily compared to CA. To improve interpretability, we changed the way we calculate EAAs for all clocks in this study (see Steps 1a-3a in Section 2.1.4). Now, we fit linear regression only on the control or baseline class (for this study, this was the samples classed as “healthy”) and then expect that if a clock captures the difference between classes, residuals for this class will be different from the control group.

Associations between estimated epigenetic age and chronological age were analysed using the Pearson correlation test, while the relationships between EAAs and sample characteristics were assessed using the Spearman correlation test, which is suitable for both continuous and ordinal variables. Two-sample *t*-tests were performed to analyse the difference in EAAs between different tissue types. All graphs presented in this study were produced using `ggplot` and its extensions (Wickham (2011)), `pheatmap` (Kolde (2019)), and base R functions (R Core Team (2009)).

### 2.2 Classifier

#### 2.2.1 Data selection

Ten different datasets spanning 990 samples were used to build the classifier. 328 were normal and 662 were healthy colon tissue samples. The classifier was trained on sex and on the epigenetic age acceleration scores from 11 different clocks.

The data was split into training and testing datasets. The training dataset consisted of data from six studies (NCBI GEO datasets GSE101764, GSE132804 450k, GSE132804 EPIC, GSE142257, GSE149282, and GSE166212), and contained 341/215 healthy/normal samples. The testing dataset included data from four studies (ArrayExpress deposited E-MTAB-3027 and E-MTAB-7036, as well as NCBI GEO datasets GSE151732 and GSE199057), and contained 321/113 healthy/normal samples. Samples originating from the same dataset were not split between training and testing sets in order to avoid potential data leakage through batch effect. The distribution of healthy and normal samples across the different datasets is provided in Table S2.

Only normal and healthy tissue samples were included when making the classifier (tumour and adenoma samples were excluded). Samples were excluded if there was no corresponding raw data (.idat) file or technical information (array identifiers and position of the sample in the array) available. Analysis of outliers using EAAd was done as described in Section 2.1.4 - samples were removed if they were outside of the mean ±3·SD interval in even one clock. In total, 39 samples were removed using these exclusion criteria.

#### 2.2.2 EAA calculation

To calculate EAAs for the classifier we used the following four-step procedure for each epigenetic clock:

- **Step 1b:** We regressed epigenetic age onto the chronological age for healthy samples in the training dataset using model (3).

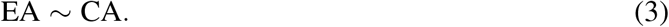
- **Step 2b:** Using linear regression coefficients obtained in Step 1b, we calculated the EAA scores for all samples used in the classifier as the regression residuals.
- **Step 3b:** We performed normalisation of the training dataset using standard normal distribution scaling.
- **Step 4b:** Test data were scaled using the mean and standard deviation of the training data used in Step 3b.

These steps were taken to prevent data leaks between the training and testing datasets. The choice of using only healthy samples in Step 1b was made to ensure a uniform EAA calculation for all epigenetic age scores, including mitotic clocks. Scaling was performed to unify the various scores’ distribution, making the classifier coefficients more interpretable. We also calculated platform-adjusted residuals by adding binary Illumina platform ID data (Illumina 450k or EPIC arrays) as a predictor in the model (3) in the first step.

#### 2.2.3 Grid search, cross-validation, and classifier training

Elastic net regression with ridge and lasso penalty terms was used when training our classifier. The optimal values for the elastic net parameters *α* and *λ* were identified through cross-validation. We manually selected folds for the cross-validation process. It was done by choosing two datasets for each fold testing data, and the remaining four for the fold training subset. By doing this, we ensured that the training and testing subsets in each fold included both healthy and normal samples, which resulted in 12 folds being used in the cross-validation process.

EAA calculation was performed separately at each fold, followed by training a classifier on the fold training set and calculating metrics on the fold testing set. This was done using a grid search for *α* ∈ [0, 1] with step 0.05, and *λ* ∈ [0, 1] with step 0.01. For each set of parameter values (fold, *α* and *λ*) we calculated two threshold-independent metrics (areas under the receiver operating characteristic (ROC-AUC) and precision-recall (PR-AUC) curves) to evaluate the model performance and identify optimal values for the parameters. For each pair of values {*α, λ*} we calculated the means of ROC-AUC across all folds and chose the optimal parameters based on the maximum mean ROC-AUC number.

The classifier model was then fitted on the training dataset using elastic net regression on EAAs and sex. The R `glmnet` (Tay et al. (2023)) and `PRROC` (Grau et al. (2015)) libraries were used to prepare the classifier and evaluate its performance metrics. Results were visualised using `pROC` (Robin et al. (2011)) and `ggplot2` (Wickham (2011)) R libraries.

## 3 RESULTS

### 3.1 Evaluation of epigenetic clocks in healthy and cancer patients

Our dataset consists of *n* = 1845 samples containing healthy (*n* = 716), normal (*n* = 522), tumour (*n* = 535), and adenoma (*n* = 72) samples from colorectal tissues (Table 2). We evaluated the relationship between chronological age and epigenetic age through Pearson correlation coefficient for each tissue category. A summary of descriptive statistics for epigenetic age scores is given in Table S3. In general, the epigenetic ages from most clocks showed positive correlations with chronological age (CA) (Figure 2A, Figure S2). In terms of correlation strength, CA and EA from first- and second-generation clocks (except Wu’s clock) have higher correlations in healthy and normal tissues (*r* = 0.46−0.79) compared to epigenetic mitotic age scores (*r <* 0.3).

**Figure 2.**
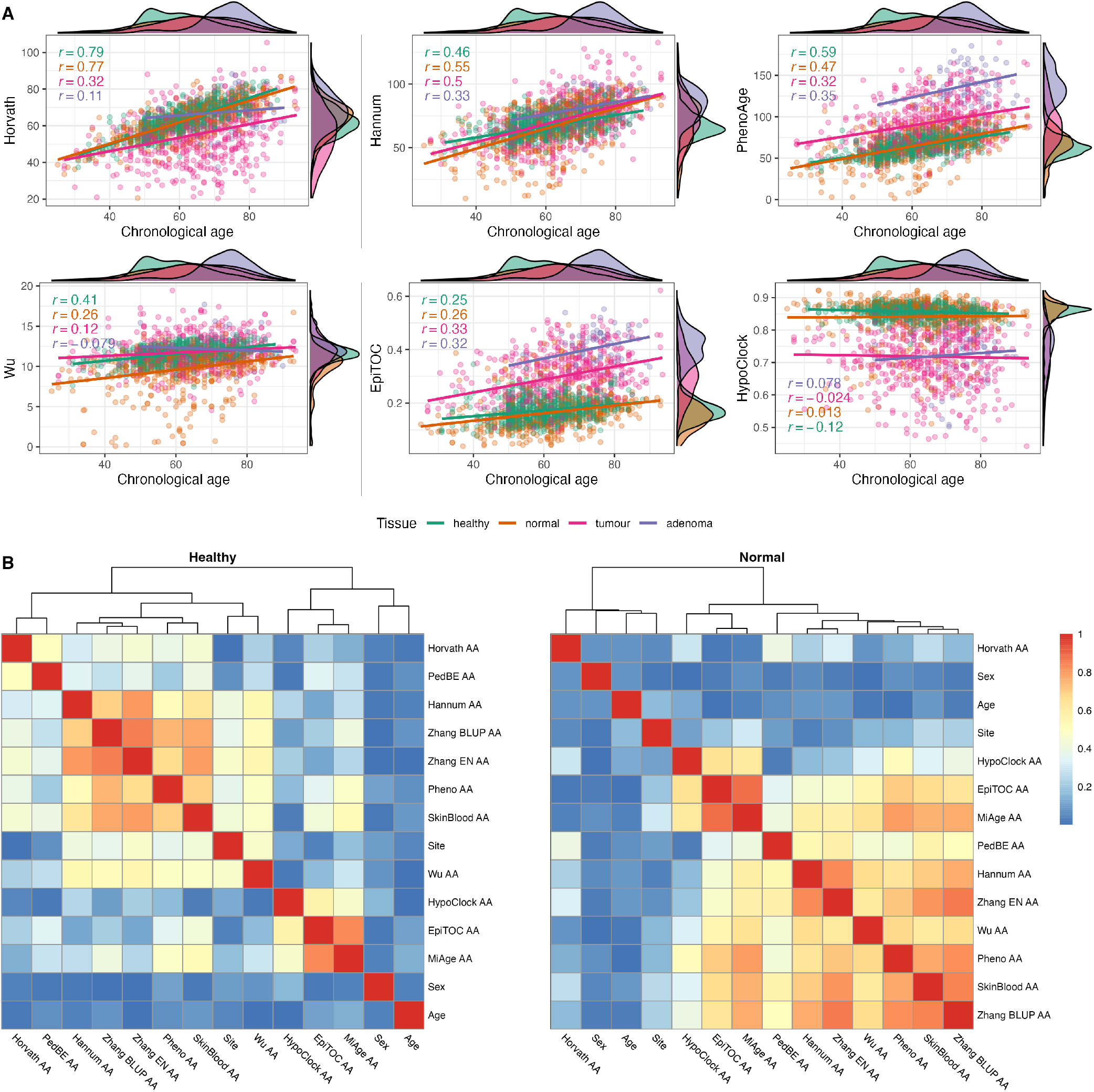
(A) Relationship between chronological age and epigenetic age estimates in four different tissues (healthy (n=716), normal (n=522), tumour (n=535), and adenoma (n=72)). Pearson’s correlation coefficients are provided for each tissue separately. (B) Heatmap of Spearman correlation (correlation coefficients are presented as absolute values) between sample characteristics and epigenetic age accelerations (EAAs) in normal colon tissues from non-CRC (healthy) and CRC (normal) participants.

We calculated EAAs following the procedure described in Section 2.1.4, the corresponding regression coefficients are given in Table S9 for Dataset 1 and Table S10 for Dataset 2. EAAs were calculated as the regression onto both CA and sex in order to reduce possible age- and sex-related bias. We analysed the relationship between EAAs and sample characteristics using the Spearman correlation test. We only included sample characteristics which were covered in more than half of the samples (i.e., age, sex, site). In all tissue samples, the correlation coefficients between EAAs and age are close to zero apart from a few EAAs from adenoma samples (Figure 2B, Figure S5), similar results were observed between EAAs and sex. On the other hand, the site (i.e., left or right colon) has a high correlation with Hannum AA and most second-generation EAAs in healthy samples, but the correlation strength is decreased in samples from CRC patients. In terms of EAAs, the first- and second-generation clock EAAs are clustered together in all tissues except for Horvath AA, PedBE AA, and Wu AA. The latter three EAAs behaved differently in CRC patients and patients with colorectal adenoma. Epigenetic mitotic clocks-based EAAs showed associations with each other, yet the coefficient became smaller in adenoma tissues (Figure S5). Analysis of unadjusted EAAs showed similar results (Figure S6). Density plots of EAA distribution in four different tissue types are given in Figure 3C and Figure S3. Summaries of EAA descriptive statistics for Dataset 1 and Dataset 2 are given in Table S4, Table S5, and Table S6.

**Figure 3.**
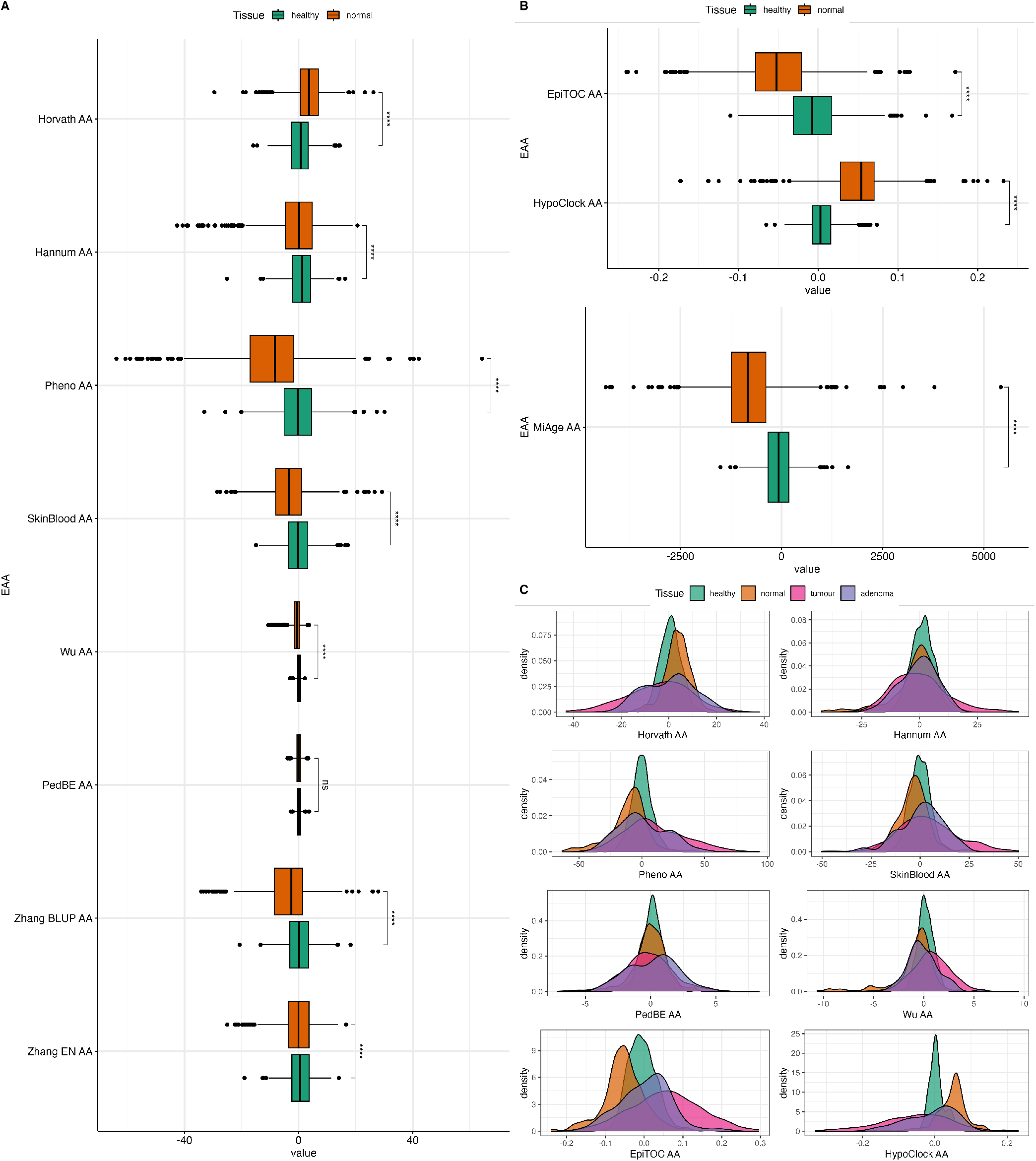
(A) Boxplots of EAAs from first- and second-generation clocks in normal colon tissues from Dataset 1. (B) Boxplots of EAAs from mitotic clocks in normal colon tissues from Dataset 1. (C) Density plots of EAA distribution in four different tissues. p-values for (A) and (B) were obtained from Welch’s two-sample t-test. ns=non significant, *p≤0.05, ** p*<*0.001, ***p*<*0.001, ****p*<*0.0001

### 3.2 Differences between EAAs in healthy individuals and CRC patients

In order to evaluate the association between epigenetic clocks and CRC, we investigated whether EAAs can capture the differences between tissues with different origins (i.e., healthy, normal, tumour, and adenoma) using the two-sample *t*-test. Among the different tissue types, tumour samples have the highest EAA variability. We also observed that Horvath AA, Pheno AA, Wu AA, EpiTOC AA, HypoClock AA, and MiAge AA captured differences between every tissue, except for healthy and adenoma (Figure S7). Interestingly, most EAAs showed significant differences between normal and adenoma samples (Figure S7). All EAAs were significantly different between normal and healthy samples, except for PedBE AA (Figure 3A, Figure 3B). Most EAAs also captured the differences between tumour and normal samples, as well as between tumour and healthy samples (Figure S7).

We repeated this test using Dataset 2 to further investigate the ability of EAAs from different epigenetic clocks to distinguishing between healthy and normal colon tissues. The distribution of EAAs from this dataset is given in Figure S4. EAAs were obtained from the residuals of regressing EA onto the CA for healthy samples and adjusted for the dataset and patient ID in Dataset 2, which contains fewer samples compared to Dataset 1. Hence, the EAA estimates will be different from the scores in the previous dataset. In general, normal samples had significantly lower EAAs compared to healthy samples. These differences were observed in all EAAs except for Horvath AA and SkinBlood AA (Figure 4). However, the p-value of SkinBlood AA was around the borderline (*p* = 0.056, 95% CI = -0.014, 1.180), hence, we may still consider SkinBlood AA for distinguishing between normal colon tissues from patients with and without CRC. This result slightly differs from comparing healthy and normal samples in the previous dataset, where PedBE AA was the only EAA that did not capture the difference between these tissues. Thus, all EAAs in our study, except for PedBE AA and Horvath AA, showed potential in discriminating between healthy and normal colon tissues in our datasets.

**Figure 4.**
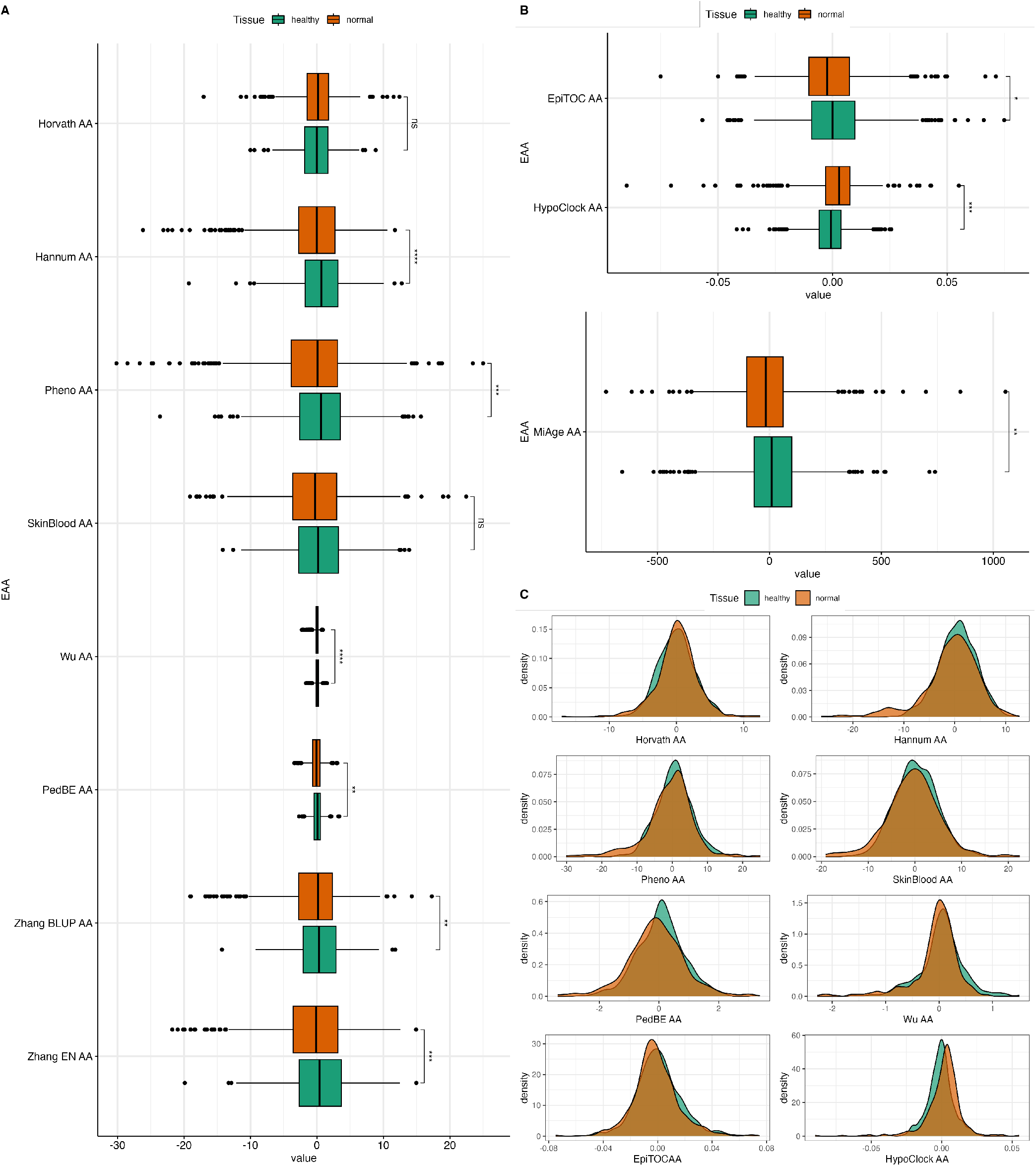
(A) Boxplots of EAAs from first- and second-generation clocks in normal colon tissues from Dataset 2. (B) Boxplots of EAAs from epigenetic clocks in normal colon tissues from Dataset 2. (C) Density plots of EAA distribution in two different tissues. The p-values were obtained from Welch’s two-sample t-test. *p≤0.05, ** p*<*0.001, ***p*<*0.001, ****p*<*0.0001.

### 3.3 EAA-based classifier demonstrates good diagnostic potential

We calculated EAAs following the steps described in Section 2.2.2, the corresponding regression coefficients and scaling parameters are given in Table S11. We trained a classifier model based on the sex data as well as on the EAAs calculated from normal colon tissue samples from six datasets, using elastic net regression with parameters *α* = 0.05 and *λ* = 0.16 estimated through the 12-folds cross-validation process (see Table S12 for the cross-validation folds list). Optimal parameter values were chosen based on the highest mean of the ROC-AUC metric across twelve cross-validation folds; heatmaps of the mean and standard deviations of the ROC-AUC are given in Figure S12. For these values of *α* and *λ*, the model selected binary sex data and ten EAAs, and excluded only Horvath’s EAA. The resulting classifier coefficients and performance were assessed on the testing subset (Table S13) and demonstrated ROC-AUC = 0.886, 95% CI [0.850, 0.922]. The ROC and PR curves for the classifier performance on the testing dataset and the histogram of the classifier’s scores are given in Figure 5A-C and Figure S10, respectively.

**Figure 5.**
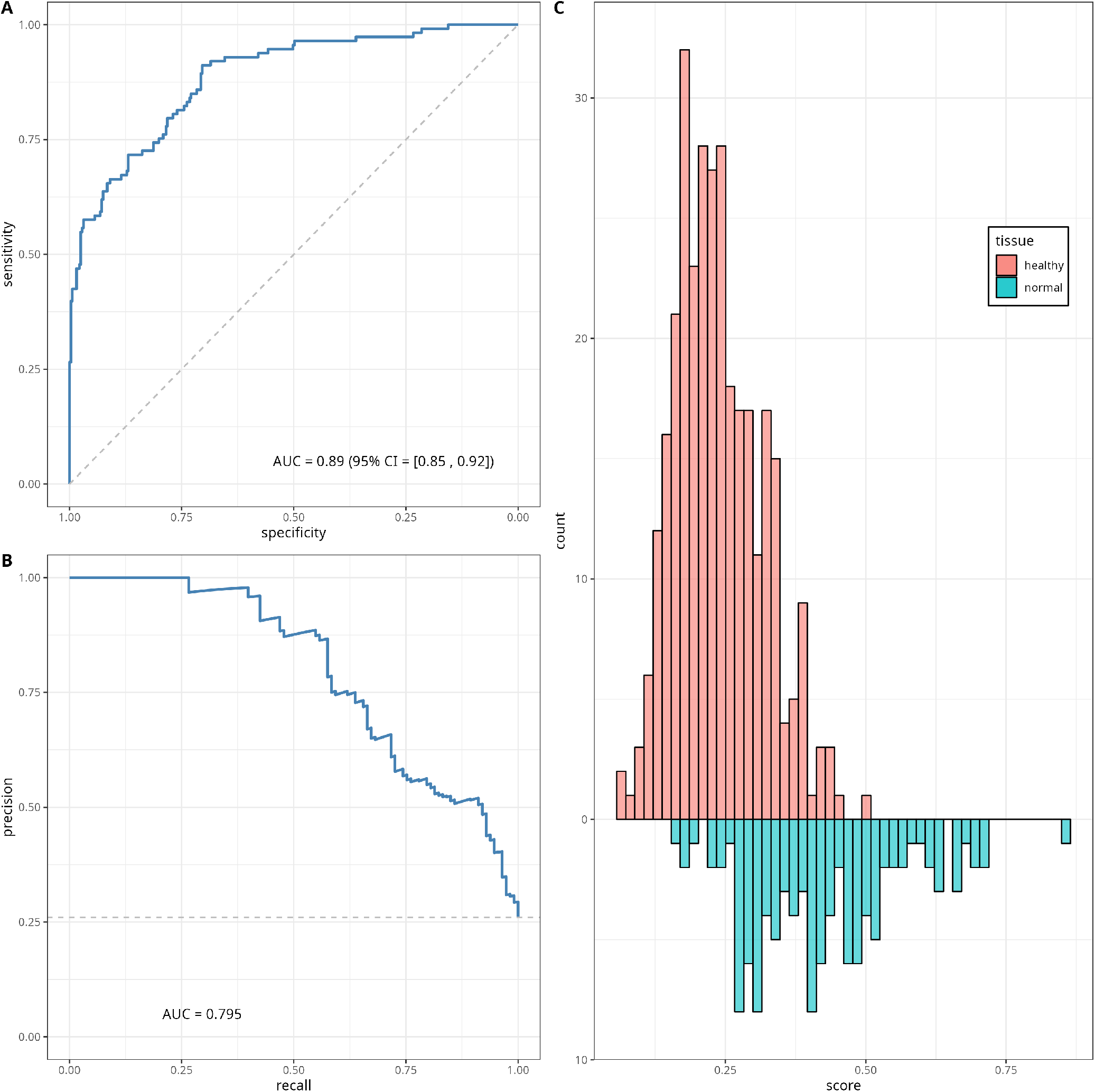
Classifier performance. ROC curve (A), precision-recall (PR) curve (B) and histogram (C) of the classifier scores for the testing data subset. The diagonal dashed line on panel (A) corresponds to the *y* = *x*, and represents the ROC of a random classifier. The horizontal line on panel (B) corresponds to the minimum precision value *y* = 0.26.

We also tried other values of the elastic net regression parameters *α* and *λ*, which have also demonstrated high values of mean ROC-AUC in the cross-validation step. In particular, for *α* = *λ* = 0.25 and *α* = 0.1, *λ* = 0.35, the classifier model used sex and six EAAs as predictors and demonstrated ROC-AUC of 0.882 (95% CI [0.845, 0.918]) and 0.835 (95% CI [0.791, 0.879]) on the testing data, respectively. The corresponding classifier coefficients for these values of regularisation parameters are presented in Table S13.

By using the EAAs adjusted for the Illumina platform ID (450k or EPIC), we trained a platform-dependent classifier. In this case, the cross-validation step was based on six folds (Table S12), and the optimal elastic net parameters values were identified as *α* = 0.05 and *λ* = 0.68. This classifier demonstrated a higher ROC-AUC=0.921 (95% CI [0.892, 0.949]) than the platform-independent version, and was based on sex and ten EAAs. The corresponding plots and coefficients can be found in Figure S11 and Table S13.

## 4 DISCUSSION

### 4.1 Associations between epigenetic age and CRC

Abnormal changes in biological age, including epigenetic age, might reflect the underlying process of cancer development, including in CRC. In our study, we focused on evaluating the relationship between epigenetic clock measures (EA and EAA) and colon tissues from participants with and without CRC. We observed that most first- and second-generation epigenetic clocks reflect the chronological age very well in normal and healthy colon tissues, especially Horvath age. On the other hand, epigenetic mitotic clocks showed weaker correlations with CA. Our results align with findings from Wang et al. (2020) and Joo et al. (2021), where Horvath and EpiTOC were reported to have the strongest and weakest associations with CA, respectively. This is not surprising, since Horvath’s clock model was originally trained to predict CA across various tissues (Horvath (2013)) while mitotic clock models were developed to account for stem cell division rates, which may affect their ability to predict CA (Yang et al. (2016)). For example, MiAge gives an estimate of cell cycle numbers (which are measured in thousands) and EpiTOC’s scores reflect the average DNAm increase due to presumed cell replication error (ranging between 0 and 1).

It is worth mentioning that associations between EA and CA vary for some of the considered clocks in histologically normal, adenoma, and cancerous colon tissues. Similar results were also described in Joo et al. (2021) for Horvath, Hannum, PhenoAge, and EpiTOC. As reviewed by Weisenberger et al. (2018), abnormal DNA methylation patterns have been observed in cancer cells, including in CRC cases. This aberration mainly results in the silencing of genes that contribute to DNA repair and tumour suppression, such as *MLH1, CDKN2A*, and *SFRP2*, hence promoting cancer growth and survival (Weisenberger et al. (2018); Schmitt and Greten (2021)). This might be a plausible explanation for the increased variance in the epigenetic age of CRC tumours. We also observed a higher variance in adenoma samples compared to normal and healthy tissues. A previous study reported that adenoma may have a similar methylation pattern with either normal colon tissue or chromosomally unstable cancer tissue, depending on the methylator epigenotype status (low or high) (Luo et al. (2014)). The variance in our data might be present due to abnormal DNAm patterns or other epigenetic instability. However, it might also be caused by the low number of adenoma samples available in this study compared to other tissues.

In general, EAAs in this study are independent of age and sex both before and after adjusting for sex, while the sample collection site correlated with some of the EAAs in healthy samples. This might be explained by the balanced ratio between male and female subjects in our dataset. Besides, evidence for sexual dimorphism in CRC is still lacking (White et al. (2018); Abancens et al. (2020)), although worldwide statistics showed slightly higher CRC incidence in males (Sung et al. (2021)). In contrast, immunological landscape variations and differentially methylated loci between the left and right colon have been observed in previous studies, which might be due to differences in the embryological lineage between the left and right colon (Kaz et al. (2014); Zhang et al. (2018); Illingworth et al. (2008)). Some CRC cases might also have higher CIMP on one side of the colon (Weisenberger et al. (2018)) and the methylated region might overlap with some of the clocks’ CpGs. However, despite the evidence, it is noteworthy that site information is available only for about half of the samples in our dataset and is distributed differently in each tissue. Hence, an explanation for the association between site and epigenetic clocks cannot be given through our study.

Our dataset consists of colon tissue with different tissue states to assess the ability of EAAs to capture the epigenetic deviation between each tissue. We observed that Pheno AA, Wu AA, and epigenetic mitotic clocks-based EAAs distinguished most of these tissues very well, compared to other EAAs. Moreover, all of the considered EAAs (except Horvath and PedBE AA) were significantly different between the healthy and normal colon tissue in both datasets. Our results are in line with Joo et al. (2021), in which EpiTOC performed well in distinguishing between these colon tissues, whereas non-mitotic clocks, especially Horvath AA, demonstrated inconsistent results. Field cancerisation that affects genomic stability, particularly the DNAm pattern, of normal colon tissues adjacent to CRC tumours might contribute to the EAA differences (Sanz-Pamplona et al. (2014)). Wang et al. (2020) also reported that normal colon tissue samples from CRC patients are differently methylated in 5-20 CpGs that overlap with CpGs from Hannum, Horvath, PhenoAge, and EpiTOC model, compared to colon tissue from participants without CRC. Hence, this might explain the sensitivity of these clocks in distinguishing normal colon tissues from individuals with different CRC diagnoses. Further investigation of the epigenome of normal colon tissue and its association with various epigenetic clock models is needed to find the most suitable CpGs as biomarkers in normal colon tissue.

### 4.2 Classifier for capturing CRC risk from normal colon tissue

The main idea behind developing a classifier was an attempt to combine the abilities of several clocks to distinguish between normal colon tissue from individuals with and without CRC. To the best of our knowledge, this is the first effort to make a cancer status predictor based on EAAs in histologically normal tissues. We performed a thorough literature search and did not manage to find any similar studies, although there were several fairly successful attempts to create CRC diagnostic methods based on peripheral blood, stool blood, and colon tissue, which are well-summarised in the recent review on CRC diagnostic, prognostic and predictive DNAm biomarkers (Mueller and Győrffy (2022)).

Our classifier demonstrated a very encouraging performance (ROC-AUC above 0.88), which is a clear indication of its diagnostic potential. The only EAA excluded from the regression by the elastic net (for *α* = 0.05, *λ* = 0.16) was Horvath AA, which is in line with the results reported in Section 3.2 and is discussed above, where Horvath EAAs were found to be distributed similarly in healthy and normal samples. At the same time, we observed that the highest absolute classifier coefficients come from EAAs derived from the Wu and PhenoAge clocks, whilst the lowest values were observed for EpiTOC, Zhang BLUP, and Skin and Blood clocks, which mostly reflects our association analyses outcomes. The improved performance of the platform-dependent classifier (ROC-AUC above 0.92) suggests that the classifier could be upgraded further with the inclusion of relevant predictors, which was not possible in the present study due to data availability. In particular, we expect that adding relevant information such as the sample location and patient ethnicity/race to the regression model could make a substantial contribution to the classifier performance. The presented framework for classifier development, including EAA calculation, cross-validation, and parameter tuning steps, could be applied to an extended (or modified) list of epigenetic clocks and relevant phenotypic data. It might also be adapted for a classifier based on DNAm data for a subset of CpGs (e.g. CpGs used in epigenetic clocks). Potentially these lead to the creation of a tool that can support diagnostic/prognostic decisions for clinical professionals.

### 4.3 Study limitations

The results presented in this paper should be considered while taking into account several shortcomings. The analysed dataset comprises data obtained from multiple independent studies which were conducted in different countries; following diverse sample extraction, processing, and storage protocols; and using four different DNAm profiling technologies (two versions of Illumina 450k and two versions of EPIC arrays). The diversity in sample handling makes our dataset very prone to technical bias. In order to reduce the influence of this bias, where possible, we pre-processed the data using consistent unified techniques and methods designed to treat samples without the context of the dataset (e.g. using single sample normalisation method `ssNoob`). We would like to point out that the heterogeneity of our data due to technical variability can be viewed as an advantage rather than as a shortcoming, since it reflects real-world data diversity.

Furthermore, the datasets from most studies had very limited clinical data available, which reduced our ability to account for several important characteristics that are known to be reflected in DNAm data. For example, sample location (i.e., left/right colon) and race are known to be associated with different distributions of EAAs (Devall et al. (2021, 2022)), which, in turn, could influence epigenetic age scores for some clocks. Hence, we cannot fully guarantee that these clocks correlate with CRC status in our dataset. Moreover, due to the limited availability of clinical data, we could not study whether the classifier scores are associated with the disease stage and outcome. This also means that when developing our model we were unable to account for some potentially important characteristics (e.g. site, cancer stage). The better performance of the platform-dependent classifier compared to the platform-independent version demonstrated that variability in the DNAm profiling platforms (Illumina arrays) influences DNAm measures and that our results could be substantially improved with a larger, more homogeneous, and better-annotated dataset.

## 5 CONCLUSION

This open access-enabled study investigated the associations between eleven epigenetic age measures and the colon tissue of individuals with and without CRC. Our results indicate that CRC status might affect the association between epigenetic age and chronological age, as well as between colon tissue EAAs and clinical characteristics. We have also demonstrated that most EAAs, except for Horvath and PedBE AA, are able to distinguish between colon tissue with different CRC status, particularly between normal and healthy colon tissues. We developed a CRC status classifier based on sex and EAAs calculated using histologically normal colon tissue DNAm data, which performed well. Although further studies on a larger, more homogeneous, and more clinically described datasets are needed to acquire a deeper understanding of this association, our results provide valuable insights into the relationship between epigenetic age and CRC. In addition, our framework could be used for developing a more robust classifier.

## AUTHOR CONTRIBUTIONS

TW, JS and OC designed the study and drafted the manuscript with input from all the authors. KP, EC, NH and VV were involved in data processing and contributed analyses. SB, VV and OC supervised the study. All authors read and approved the final version of the manuscript.

## ABBREVIATIONS

AA: Age acceleration
AUC: Area under the curve
BLUP: Best linear unbiased prediction
CA: Chronological age
CIMP: CpG island methylator phenotype
CpG: Cytosine-phosphate-Guanine
CRC: Colorectal cancer
DNAm: DNA methylation
EA: Epigenetic age
EAA: Epigenetic age acceleration
EMBL-EBI: European Molecular Biology Laboratory-European Bioinformatics Institute
EN: Elastic net
NCBI GEO: National Center for Biotechnology Information - Gene Expression Omnibus
PCGT: Polycomb group target
PedBE: Pediatric-Buccal-Epigenetic
PR: Precision-Recall
ROC: Receiver Operating Characteristic

## ACKNOWLEDGEMENTS

The authors are grateful to the studies which made their data openly available. We also thank the UCL Cancer Institute Medical Genomics lab for the stimulating and inspiring discussions.

## SUPPLEMENTAL DATA

Supplementary Material should be uploaded separately on submission, if there are Supplementary Figures, please include the caption in the same file as the figure. LaTeX Supplementary Material templates can be found in the Frontiers LaTeX folder.

## DATA AVAILABILITY STATEMENT

The datasets used for this study are openly available from NCBI GEO and EMBL-EBI ArrayExpress repositories using unique accession IDs. The list of the accession number(s) can be found in Table S1. A copy of the table with clinical data and calculated epigenetic age together with the code is openly available from the UCL Medical Genomics Lab GitHub repository.

