## supplementary tables and figures for "Open access-enabled evaluation of epigenetic age acceleration in colorectal cancer and development of a classifier with diagnostic potential"

### Supplementary Material

#### 1 SUPPLEMENTARY TABLES AND FIGURES

##### 1.1 Supplementary Tables

**Table S1.** List of datasets used in this study and sample distribution in each dataset.

| Dataset ID | Platform | Dataset 1 | | | | Dataset 2 | |
|---|---|---|---|---|---|---|---|
| | | Healthy | Normal | Tumour | Adenoma | Healthy | Normal |
| E-MTAB-7036 | 450K | 0 | 27 | 189 | 0 | 0 | 27 |
| E-MTAB-7854 | EPIC | 0 | 0 | 0 | 64 | 0 | 0 |
| E-MTAB-3027 | 450K | 0 | 23 | 23 | 0 | 0 | 23 |
| GSE101764 | 450K | 0 | 144 | 103 | 0 | 0 | 144 |
| GSE131013 | 450K | 48 | 96 | 93 | 0 | 48 | 94 |
| GSE132804 | 450K | 73 | 20 | 0 | 0 | 73 | 20 |
| GSE132804 | EPIC | 151 | 49 | 0 | 0 | 151 | 46 |
| GSE142257 | EPIC | 118 | 0 | 0 | 0 | 118 | 0 |
| GSE149282 | EPIC | 0 | 10 | 10 | 0 | 0 | 9 |
| GSE151732 | EPIC | 254 | 0 | 0 | 0 | 253 | 0 |
| GSE159898 | EPIC | 0 | 16 | 22 | 0 | 0 | 14 |
| GSE166212 | EPIC | 0 | 6 | 24 | 8 | 0 | 5 |
| GSE171550 | EPIC | 0 | 52 | 0 | 0 | 0 | 47 |
| GSE199057 | EPIC | 72 | 79 | 71 | 0 | 72 | 76 |

**Table S2.** Sample distribution in datasets used for classifier.

| Dataset | Healthy | Normal | Platform | Test/Train |
|---|---|---|---|---|
| E-MTAB-7036 | 0 | 24 | 450k | Test |
| E-MTAB-3027 | 0 | 22 | 450k | Test |
| GSE101764 | 0 | 140 | 450k | Train |
| GSE132804 | 73 | 17 | 450k | Train |
| GSE132804 | 150 | 44 | EPIC | Train |
| GSE142257 | 118 | 0 | EPIC | Train |
| GSE149282 | 0 | 9 | EPIC | Train |
| GSE151732 | 250 | 0 | EPIC | Test |
| GSE166212 | 0 | 5 | EPIC | Train |
| GSE199057 | 71 | 67 | EPIC | Test |

**Table S3.** Summary of epigenetic age (EA) data. p-values obtained from Welch's t-test, testing the difference between male and female groups for each EA ($H0$: mean value of the EA is the same for male and female groups). Significant differences ($p < 0.05$) are highlighted with bold text.

| Clock | Tissue | Mean (SD) | | | p-value | 95% CI |
|---|---|---|---|---|---|---|
| | | All | Female | Male | | |
| **Horvath** | Healthy | 62.12 (7.74) | 61.55 (7.56) | 63.1 (7.97) | **0.011** | (-2.75, -0.36) |
| | Normal | 63.61 (9.58) | 62.86 (9.57) | 64.1 (9.57) | 0.148 | (-2.92, 0.44) |
| | Tumour | 55.48 (14.14) | 54.91 (14.84) | 55.91 (13.6) | 0.427 | (-3.45, 1.46) |
| | Adenoma | 67.6 (9.91) | 68.71 (9.35) | 65.36 (10.81) | 0.202 | (-1.87, 8.58) |
| **Hannum** | Healthy | 66.63 (9.19) | 65.92 (8.95) | 67.86 (9.49) | **0.007** | (-3.35, -0.52) |
| | Normal | 67.28 (18.35) | 65.94 (18.24) | 68.15 (18.4) | 0.178 | (-5.44, 1.01) |
| | Tumour | 72.45 (17.77) | 73.73 (18.7) | 71.49 (17.02) | 0.155 | (-0.85, 5.33) |
| | Adenoma | 83.04 (11.45) | 84.43 (9.41) | 80.24 (14.55) | 0.208 | (-2.45, 10.84) |
| **PhenoAge** | Healthy | 62.84 (10.41) | 61.84 (10.48) | 64.57 (10.06) | **0.001** | (-4.28, -1.17) |
| | Normal | 66.34 (20.3) | 65.57 (21.06) | 66.84 (19.81) | 0.491 | (-4.89, 2.35) |
| | Tumour | 92.93 (26.37) | 95.05 (29.19) | 91.35 (23.97) | 0.119 | (-0.96, 8.34) |
| | Adenoma | 136.16 (21.11) | 137.17 (18.93) | 134.13 (25.24) | 0.606 | (-8.79, 14.86) |
| **SkinBlood** | Healthy | 70.82 (8.63) | 70.73 (8.7) | 70.98 (8.51) | 0.715 | (-1.55, 1.06) |
| | Normal | 69.61 (12.29) | 69.14 (13.06) | 69.92 (11.77) | 0.486 | (-3, 1.43) |
| | Tumour | 81.26 (18.32) | 82.6 (19.32) | 80.26 (17.5) | 0.15 | (-0.85, 5.52) |
| | Adenoma | 102.44 (11.49) | 104.74 (10.79) | 97.82 (11.66) | **0.019** | (1.18, 12.66) |
| **PedBE** | Healthy | 13.72 (1.71) | 13.68 (1.69) | 13.79 (1.76) | 0.422 | (-0.37, 0.16) |
| | Normal | 12.71 (2.1) | 12.69 (2.1) | 12.73 (2.11) | 0.846 | (-0.41, 0.33) |
| | Tumour | 12.47 (2.8) | 12.35 (2.92) | 12.55 (2.71) | 0.424 | (-0.68, 0.29) |
| | Adenoma | 15.21 (2.48) | 15.28 (2.22) | 15.07 (2.97) | 0.767 | (-1.19, 1.6) |
| **Wu** | Healthy | 11.53 (1) | 11.47 (0.97) | 11.65 (1.03) | **0.021** | (-0.33, -0.03) |
| | Normal | 9.73 (2.53) | 9.65 (2.43) | 9.78 (2.59) | 0.571 | (-0.57, 0.31) |
| | Tumour | 11.82 (2.02) | 11.71 (1.87) | 11.91 (2.12) | 0.23 | (-0.55, 0.13) |
| | Adenoma | 12.11 (1.63) | 11.86 (1.25) | 12.6 (2.14) | 0.124 | (-1.71, 0.22) |
| **Zhang BLUP** | Healthy | 73.81 (7.68) | 73.47 (7.76) | 74.4 (7.53) | 0.115 | (-2.09, 0.23) |
| | Normal | 71.32 (12.38) | 71.17 (13.07) | 71.41 (11.93) | 0.829 | (-2.47, 1.98) |
| | Tumour | 82.54 (15.55) | 84.14 (16.76) | 81.34 (14.5) | **0.044** | (0.08, 5.52) |
| | Adenoma | 109.19 (9.71) | 108.48 (8.72) | 110.6 (11.52) | 0.432 | (-7.53, 3.29) |
| **Zhang EN** | Healthy | 75.82 (7.45) | 75.46 (7.52) | 76.44 (7.31) | 0.085 | (-2.11, 0.14) |
| | Normal | 74.37 (10.83) | 73.84 (11.12) | 74.72 (10.64) | 0.368 | (-2.81, 1.04) |
| | Tumour | 76.67 (11.69) | 77.54 (12.39) | 76.01 (11.11) | 0.141 | (-0.51, 3.56) |
| | Adenoma | 98.61 (7.33) | 99.24 (6.53) | 97.37 (8.73) | 0.36 | (-2.22, 5.96) |
| **EpiTOC** | Healthy | 0.17 (0.04) | 0.17 (0.04) | 0.17 (0.04) | 0.954 | (-0.01, 0.01) |
| | Normal | 0.17 (0.07) | 0.16 (0.07) | 0.17 (0.07) | 0.534 | (-0.02, 0.01) |
| | Tumour | 0.3 (0.09) | 0.31 (0.1) | 0.29 (0.08) | **0.003** | (0.01, 0.04) |
| | Adenoma | 0.4 (0.07) | 0.41 (0.06) | 0.39 (0.08) | 0.376 | (-0.02, 0.05) |
| **HypoClock** | Healthy | 0.86 (0.02) | 0.86 (0.02) | 0.86 (0.02) | **0.035** | (0, 0.01) |
| | Normal | 0.84 (0.06) | 0.84 (0.06) | 0.84 (0.07) | 0.33 | (-0.01, 0.02) |
| | Tumour | 0.72 (0.09) | 0.72 (0.09) | 0.72 (0.09) | 0.568 | (-0.02, 0.01) |
| | Adenoma | 0.72 (0.08) | 0.74 (0.07) | 0.69 (0.08) | **0.01** | (0.01, 0.09) |
| **MiAge** | Healthy | 1990.07 (512.67) | 1998.84 (492.43) | 1974.98 (546.42) | 0.56 | (-56.45, 104.16) |
| | Normal | 1708.7 (1032.78) | 1691.57 (1129.02) | 1719.87 (966.6) | 0.767 | (-216.3, 159.72) |
| | Tumour | 3942.91 (1747.98) | 4232.13 (1832.32) | 3726.46 (1652.25) | **0.001** | (203.9, 807.45) |
| | Adenoma | 5484.91 (1522.72) | 5339.43 (1437.43) | 5775.87 (1674.18) | 0.281 | (-1244.2, 371.32) |

**Table S4.** Summary of epigenetic age acceleration (EAA) scores (sex-adjusted) from Dataset 1. p-values obtained from Welch's t-test, testing the difference between male and female groups for each EAA ($H0$: mean value of the EAA is the same for male and female groups). Significant differences ($p < 0.05$) are highlighted with bold text.

| EAA | Tissue | Mean (SD) | | | p-value | 95% CI |
|---|---|---|---|---|---|---|
| | | All | Female | Male | | |
| **Horvath AA** | Healthy | 0.593 (4.351) | 0.691 (4.186) | 0.423 (4.625) | 0.439 | (-0.412, 0.949) |
| | Normal | 3.481 (5.84) | 3.91 (5.844) | 3.201 (5.829) | 0.176 | (-0.319, 1.736) |
| | Tumour | -4.19 (13.488) | -4.17 (14.321) | -4.206 (12.853) | 0.976 | (-2.318, 2.391) |
| | Adenoma | 0.006 (10.224) | -0.654 (9.966) | 1.327 (10.816) | 0.456 | (-7.296, 3.333) |
| **Hannum AA** | Healthy | 1.077 (4.865) | 1.134 (4.962) | 0.978 (4.7) | 0.674 | (-0.574, 0.887) |
| | Normal | -1.39 (10.057) | -0.771 (9.96) | -1.793 (10.115) | 0.256 | (-0.742, 2.785) |
| | Tumour | -0.014 (11.892) | 2.278 (11.928) | -1.729 (11.59) | **<0.001** | (1.985, 6.031) |
| | Adenoma | -0.53 (8.013) | -0.295 (7.376) | -1.001 (9.311) | 0.747 | (-3.704, 5.117) |
| **Pheno AA** | Healthy | -0.169 (7.902) | 0.56 (7.795) | -1.424 (7.943) | **0.001** | (0.782, 3.185) |
| | Normal | -10.45 (15.836) | -9.46 (15.751) | -11.095 (15.882) | 0.248 | (-1.145, 4.417) |
| | Tumour | 10.327 (24.186) | 12.405 (26.268) | 8.771 (22.421) | 0.093 | (-0.607, 7.874) |
| | Adenoma | 0.71 (18.79) | -0.518 (19.028) | 3.164 (18.455) | 0.434 | (-13.059, 5.694) |
| **SkinBlood AA** | Healthy | -0.178 (5.188) | -0.215 (5.297) | -0.114 (5.002) | 0.798 | (-0.88, 0.677) |
| | Normal | -3.661 (7.933) | -3.928 (8.158) | -3.486 (7.791) | 0.538 | (-1.853, 0.968) |
| | Tumour | 3.649 (15.474) | 3.892 (16.3) | 3.467 (14.849) | 0.757 | (-2.27, 3.12) |
| | Adenoma | 1.194 (10.834) | 1.437 (11.262) | 0.708 (10.139) | 0.783 | (-4.555, 6.013) |
| **PedBE AA** | Healthy | 0.126 (0.85) | 0.112 (0.837) | 0.149 (0.873) | 0.572 | (-0.169, 0.093) |
| | Normal | 0.022 (1.209) | 0.07 (1.217) | -0.01 (1.205) | 0.465 | (-0.134, 0.293) |
| | Tumour | -0.213 (1.948) | -0.325 (2.034) | -0.128 (1.881) | 0.253 | (-0.535, 0.141) |
| | Adenoma | 0.174 (1.848) | 0.008 (1.751) | 0.506 (2.024) | 0.31 | (-1.476, 0.481) |
| **Wu AA** | Healthy | 0.172 (0.779) | 0.118 (0.749) | 0.264 (0.822) | **0.018** | (-0.267, -0.025) |
| | Normal | -1.02 (2.231) | -1.012 (2.188) | -1.025 (2.262) | 0.944 | (-0.376, 0.404) |
| | Tumour | 0.769 (1.934) | 0.633 (1.88) | 0.871 (1.971) | 0.157 | (-0.567, 0.092) |
| | Adenoma | -0.028 (1.588) | -0.214 (1.259) | 0.344 (2.08) | 0.236 | (-1.499, 0.384) |
| **Zhang BLUP AA** | Healthy | 0.218 (4.833) | 0.237 (4.793) | 0.185 (4.91) | 0.889 | (-0.689, 0.794) |
| | Normal | -4.053 (9.205) | -3.712 (9.27) | -4.275 (9.169) | 0.497 | (-1.062, 2.187) |
| | Tumour | 3.495 (12.305) | 4.194 (13.087) | 2.971 (11.68) | 0.263 | (-0.923, 3.371) |
| | Adenoma | 1.248 (9.155) | -0.766 (8.104) | 5.278 (9.955) | **0.014** | (-10.788, -1.3) |
| **Zhang EN AA** | Healthy | 0.624 (4.462) | 0.644 (4.501) | 0.591 (4.404) | 0.878 | (-0.623, 0.729) |
| | Normal | -0.784 (6.521) | -0.564 (6.577) | -0.927 (6.491) | 0.537 | (-0.789, 1.514) |
| | Tumour | -0.208 (8.422) | 0.444 (8.698) | -0.696 (8.19) | 0.125 | (-0.317, 2.597) |
| | Adenoma | 1.019 (6.152) | 0.201 (5.713) | 2.655 (6.777) | 0.135 | (-5.709, 0.801) |
| **EpiTOC AA** | Healthy | -0.005 (0.036) | -0.005 (0.034) | -0.005 (0.039) | 0.96 | (-0.006, 0.006) |
| | Normal | -0.051 (0.054) | -0.054 (0.055) | -0.049 (0.053) | 0.321 | (-0.014, 0.005) |
| | Tumour | 0.056 (0.088) | 0.062 (0.092) | 0.052 (0.085) | 0.179 | (-0.005, 0.026) |
| | Adenoma | 0.004 (0.064) | 0.002 (0.059) | 0.007 (0.075) | 0.814 | (-0.039, 0.031) |
| **HypoClock AA** | Healthy | 0.006 (0.021) | 0.003 (0.019) | 0.011 (0.024) | **<0.001** | (-0.011, -0.004) |
| | Normal | 0.049 (0.045) | 0.051 (0.046) | 0.047 (0.045) | 0.329 | (-0.004, 0.012) |
| | Tumour | -0.054 (0.092) | -0.052 (0.094) | -0.056 (0.09) | 0.694 | (-0.013, 0.019) |
| | Adenoma | -0.009 (0.065) | 0.004 (0.059) | -0.033 (0.071) | **0.036** | (0.003, 0.071) |
| **MiAge AA** | Healthy | -72.183 (399.83) | -87.704 (385.32) | -45.448 (423.077) | 0.184 | (-104.643, 20.13) |
| | Normal | -826.846 (929.711) | -878.407 (1005.168) | -793.233 (877.051) | 0.321 | (-253.595, 83.246) |
| | Tumour | 891.199 (1608.919) | 1013.286 (1627.703) | 799.833 (1591.255) | 0.13 | (-63.337, 490.243) |
| | Adenoma | 90.346 (1437.475) | -19.894 (1404.599) | 310.826 (1506.991) | 0.374 | (-1073.536, 412.097) |

**Table S5.** Summary of epigenetic age acceleration (EAA) scores (unadjusted) from Dataset 1. p-values obtained from Welch's t-test, testing the difference between male and female groups for each EAA ($H0$: mean value of the EAA is the same for male and female groups). Significant differences ($p < 0.05$) are highlighted with bold text.

| EAA | Tissue | Mean (SD) | | | p-value | 95% CI |
|---|---|---|---|---|---|---|
| | | All | Female | Male | | |
| **Horvath AA** | Healthy | 0.565 (4.336) | 0.303 (4.153) | 1.016 (4.607) | **0.039** | (-1.39, -0.036) |
| | Normal | 3.489 (5.83) | 3.251 (5.848) | 3.645 (5.822) | 0.451 | (-1.422, 0.633) |
| | Tumour | -4.163 (13.523) | -4.765 (14.351) | -3.712 (12.873) | 0.381 | (-3.412, 1.306) |
| | Adenoma | 0.018 (10.296) | -1.005 (9.986) | 2.063 (10.813) | 0.251 | (-8.385, 2.248) |
| **Hannum AA** | Healthy | 1.054 (4.869) | 0.709 (4.927) | 1.648 (4.717) | **0.012** | (-1.669, -0.209) |
| | Normal | -1.389 (10.082) | -1.48 (9.988) | -1.33 (10.158) | 0.868 | (-1.919, 1.62) |
| | Tumour | 0.015 (11.881) | 1.652 (12.001) | -1.21 (11.661) | **0.006** | (0.827, 4.898) |
| | Adenoma | -0.521 (8.115) | -0.707 (7.443) | -0.149 (9.481) | 0.802 | (-5.04, 3.925) |
| **Pheno AA** | Healthy | -0.224 (7.85) | -0.222 (7.791) | -0.227 (7.964) | 0.993 | (-1.198, 1.208) |
| | Normal | -10.438 (15.773) | -10.799 (15.688) | -10.202 (15.849) | 0.672 | (-3.37, 2.175) |
| | Tumour | 10.383 (24.092) | 11.2 (26.229) | 9.771 (22.385) | 0.508 | (-2.805, 5.663) |
| | Adenoma | 0.752 (18.921) | -1.21 (19.008) | 4.675 (18.513) | 0.214 | (-15.278, 3.509) |
| **SkinBlood AA** | Healthy | -0.173 (5.188) | -0.153 (5.298) | -0.208 (5.001) | 0.889 | (-0.723, 0.834) |
| | Normal | -3.662 (7.936) | -3.823 (8.164) | -3.557 (7.795) | 0.711 | (-1.678, 1.146) |
| | Tumour | 3.644 (15.479) | 3.987 (16.304) | 3.388 (14.853) | 0.662 | (-2.096, 3.295) |
| | Adenoma | 1.19 (10.841) | 1.492 (11.265) | 0.587 (10.146) | 0.733 | (-4.382, 6.192) |
| **PedBE AA** | Healthy | 0.125 (0.849) | 0.104 (0.836) | 0.16 (0.872) | 0.4 | (-0.187, 0.075) |
| | Normal | 0.022 (1.208) | 0.058 (1.216) | -0.002 (1.204) | 0.583 | (-0.154, 0.273) |
| | Tumour | -0.212 (1.948) | -0.336 (2.033) | -0.119 (1.88) | 0.209 | (-0.555, 0.122) |
| | Adenoma | 0.174 (1.848) | 0.001 (1.751) | 0.52 (2.022) | 0.29 | (-1.497, 0.458) |
| **Wu AA** | Healthy | 0.168 (0.785) | 0.068 (0.746) | 0.34 (0.821) | **<0.001** | (-0.393, -0.15) |
| | Normal | -1.019 (2.23) | -1.096 (2.185) | -0.969 (2.261) | 0.524 | (-0.516, 0.263) |
| | Tumour | 0.773 (1.937) | 0.558 (1.877) | 0.934 (1.969) | **0.025** | (-0.705, -0.047) |
| | Adenoma | -0.025 (1.602) | -0.259 (1.261) | 0.442 (2.082) | 0.14 | (-1.642, 0.242) |
| **Zhang BLUP AA** | Healthy | 0.205 (4.839) | 0.055 (4.794) | 0.463 (4.915) | 0.281 | (-1.149, 0.334) |
| | Normal | -4.051 (9.199) | -4.022 (9.262) | -4.07 (9.171) | 0.953 | (-1.576, 1.672) |
| | Tumour | 3.508 (12.302) | 3.918 (13.091) | 3.202 (11.688) | 0.513 | (-1.432, 2.864) |
| | Adenoma | 1.262 (9.236) | -0.928 (8.113) | 5.641 (9.939) | **0.008** | (-11.308, -1.829) |
| **Zhang EN AA** | Healthy | 0.612 (4.461) | 0.467 (4.491) | 0.862 (4.408) | 0.251 | (-1.071, 0.281) |
| | Normal | -0.782 (6.519) | -0.86 (6.575) | -0.731 (6.493) | 0.826 | (-1.28, 1.023) |
| | Tumour | -0.196 (8.429) | 0.179 (8.714) | -0.476 (8.212) | 0.378 | (-0.804, 2.116) |
| | Adenoma | 1.035 (6.205) | 0.043 (5.721) | 3.019 (6.768) | 0.072 | (-6.229, 0.277) |
| **EpiTOC AA** | Healthy | -0.005 (0.036) | -0.005 (0.034) | -0.006 (0.039) | 0.752 | (-0.005, 0.007) |
| | Normal | -0.051 (0.054) | -0.053 (0.055) | -0.049 (0.053) | 0.415 | (-0.014, 0.006) |
| | Tumour | 0.056 (0.088) | 0.063 (0.092) | 0.051 (0.085) | 0.146 | (-0.004, 0.027) |
| | Adenoma | 0.004 (0.064) | 0.003 (0.059) | 0.006 (0.075) | 0.852 | (-0.039, 0.032) |
| **HypoClock AA** | Healthy | 0.006 (0.021) | 0.004 (0.019) | 0.009 (0.024) | **0.003** | (-0.009, -0.002) |
| | Normal | 0.049 (0.045) | 0.053 (0.046) | 0.046 (0.045) | 0.098 | (-0.001, 0.015) |
| | Tumour | -0.054 (0.092) | -0.051 (0.094) | -0.057 (0.09) | 0.462 | (-0.01, 0.022) |
| | Adenoma | -0.009 (0.065) | 0.004 (0.059) | -0.035 (0.071) | **0.024** | (0.005, 0.073) |
| **MiAge AA** | Healthy | -71.386 (398.219) | -76.445 (384.038) | -62.672 (422.14) | 0.664 | (-75.999, 48.452) |
| | Normal | -827.019 (929.725) | -859.119 (1006.151) | -806.093 (877.344) | 0.537 | (-221.575, 115.524) |
| | Tumour | 890.387 (1610.832) | 1030.627 (1628.608) | 785.436 (1591.987) | 0.083 | (-31.742, 522.124) |
| | Adenoma | 89.706 (1436.211) | -9.966 (1404.747) | 289.051 (1507.596) | 0.422 | (-1042.07, 444.037) |

**Table S6.** Summary of epigenetic age acceleration (EAA) scores from Dataset 2. p-values obtained from Welch's t-test, testing the difference between male and female groups for each EAA ($H0$: mean value of the EAA is the same for male and female groups). Significant differences ($p < 0.05$) are highlighted with bold text.

| EAA | Tissue | Mean (SD) | | | p-value | 95% CI |
|---|---|---|---|---|---|---|
| | | All | Female | Male | | |
| **Sex-adjusted** | | | | | | |
| **Horvath AA** | Healthy | -0.041 (2.692) | 0.073 (2.729) | -0.237 (2.621) | 0.135 | (-0.096, 0.715) |
| | Normal | 0.058 (3.106) | 0.208 (3.014) | -0.039 (3.165) | 0.379 | (-0.304, 0.798) |
| **Hannum AA** | Healthy | 0.496 (3.806) | 0.575 (3.946) | 0.359 (3.554) | 0.453 | (-0.348, 0.78) |
| | Normal | -0.702 (5.458) | -0.011 (5.238) | -1.143 (5.558) | **0.021** | (0.17, 2.094) |
| **Pheno AA** | Healthy | 0.535 (5.048) | 0.772 (5.24) | 0.126 (4.68) | 0.089 | (-0.099, 1.393) |
| | Normal | -0.758 (6.878) | 0.144 (6.296) | -1.335 (7.177) | **0.015** | (0.285, 2.671) |
| **SkinBlood AA** | Healthy | 0.241 (4.461) | 0.17 (4.586) | 0.365 (4.242) | 0.564 | (-0.862, 0.471) |
| | Normal | -0.342 (5.712) | -0.192 (5.716) | -0.437 (5.716) | 0.638 | (-0.78, 1.27) |
| **PedBE AA** | Healthy | 0.069 (0.791) | 0.063 (0.787) | 0.079 (0.8) | 0.8 | (-0.137, 0.106) |
| | Normal | -0.098 (0.946) | -0.049 (0.889) | -0.129 (0.98) | 0.344 | (-0.086, 0.246) |
| **Wu AA** | Healthy | 0.046 (0.415) | 0.035 (0.435) | 0.065 (0.379) | 0.334 | (-0.091, 0.031) |
| | Normal | -0.066 (0.446) | -0.054 (0.439) | -0.073 (0.45) | 0.642 | (-0.061, 0.098) |
| **Zhang BLUP AA** | Healthy | 0.323 (3.554) | 0.318 (3.556) | 0.332 (3.557) | 0.96 | (-0.556, 0.528) |
| | Normal | -0.457 (4.874) | 0.089 (4.687) | -0.806 (4.966) | **0.041** | (0.035, 1.756) |
| **Zhang EN AA** | Healthy | 0.508 (4.619) | 0.557 (4.644) | 0.423 (4.584) | 0.708 | (-0.568, 0.836) |
| | Normal | -0.72 (6.011) | -0.109 (5.784) | -1.11 (6.129) | 0.065 | (-0.061, 2.063) |
| **EpiTOC AA** | Healthy | 0.001 (0.017) | 0 (0.017) | 0.002 (0.016) | 0.36 | (-0.004, 0.001) |
| | Normal | -0.001 (0.016) | -0.001 (0.015) | -0.002 (0.017) | 0.548 | (-0.002, 0.004) |
| **HypoClock AA** | Healthy | -0.001 (0.009) | -0.001 (0.009) | 0 (0.009) | 0.099 | (-0.003, 0) |
| | Normal | 0.001 (0.013) | 0 (0.012) | 0.002 (0.013) | 0.137 | (-0.004, 0.001) |
| **MiAge AA** | Healthy | 11.357 (157.03) | 3.819 (166.835) | 24.392 (137.784) | 0.076 | (-43.297, 2.15) |
| | Normal | -16.08 (166.637) | -14.158 (153.926) | -17.31 (174.516) | 0.831 | (-25.94, 32.245) |
| **Unadjusted** | | | | | | |
| **Horvath AA** | Healthy | -0.055 (2.688) | -0.116 (2.728) | 0.052 (2.618) | 0.415 | (-0.574, 0.237) |
| | Normal | 0.077 (3.108) | -0.139 (3.011) | 0.216 (3.165) | 0.206 | (-0.906, 0.196) |
| **Hannum AA** | Healthy | 0.485 (3.816) | 0.327 (3.941) | 0.757 (3.581) | 0.137 | (-0.996, 0.137) |
| | Normal | -0.686 (5.502) | -0.489 (5.303) | -0.812 (5.63) | 0.515 | (-0.652, 1.297) |
| **Pheno AA** | Healthy | 0.524 (5.074) | 0.437 (5.254) | 0.676 (4.754) | 0.534 | (-0.993, 0.515) |
| | Normal | -0.742 (7.005) | -0.517 (6.444) | -0.887 (7.348) | 0.553 | (-0.852, 1.59) |
| **SkinBlood AA** | Healthy | 0.247 (4.466) | 0.243 (4.592) | 0.254 (4.246) | 0.974 | (-0.678, 0.656) |
| | Normal | -0.35 (5.723) | -0.067 (5.725) | -0.531 (5.724) | 0.375 | (-0.563, 1.491) |
| **PedBE AA** | Healthy | 0.069 (0.791) | 0.06 (0.787) | 0.084 (0.799) | 0.697 | (-0.145, 0.097) |
| | Normal | -0.098 (0.945) | -0.055 (0.888) | -0.125 (0.98) | 0.411 | (-0.096, 0.235) |
| **Wu AA** | Healthy | 0.046 (0.416) | 0.026 (0.435) | 0.08 (0.379) | 0.079 | (-0.116, 0.006) |
| | Normal | -0.065 (0.446) | -0.073 (0.439) | -0.059 (0.451) | 0.723 | (-0.094, 0.065) |
| **Zhang BLUP AA** | Healthy | 0.317 (3.56) | 0.227 (3.558) | 0.474 (3.566) | 0.371 | (-0.791, 0.296) |
| | Normal | -0.449 (4.881) | -0.078 (4.702) | -0.686 (4.985) | 0.167 | (-0.255, 1.471) |
| **Zhang EN AA** | Healthy | 0.497 (4.623) | 0.384 (4.642) | 0.694 (4.593) | 0.387 | (-1.013, 0.393) |
| | Normal | -0.704 (6.005) | -0.4 (5.793) | -0.898 (6.138) | 0.358 | (-0.565, 1.561) |
| **EpiTOC AA** | Healthy | 0.001 (0.017) | 0.001 (0.017) | 0.001 (0.016) | 0.608 | (-0.003, 0.002) |
| | Normal | -0.001 (0.016) | 0 (0.015) | -0.002 (0.017) | 0.287 | (-0.001, 0.004) |
| **HypoClock AA** | Healthy | -0.001 (0.009) | -0.001 (0.009) | -0.001 (0.009) | 0.607 | (-0.002, 0.001) |
| | Normal | 0.001 (0.013) | 0.001 (0.013) | 0.002 (0.013) | 0.564 | (-0.003, 0.002) |
| **MiAge AA** | Healthy | 11.592 (156.914) | 6.911 (166.966) | 19.686 (137.746) | 0.27 | (-35.503, 9.954) |
| | Normal | -16.413 (166.859) | -8.101 (154.073) | -21.729 (174.585) | 0.358 | (-15.485, 42.741) |

**Table S7.** EAA differences between four different tissue (healthy ($n = 716$), normal ($n = 522$), tumour ($n = 535$), adenoma ($n = 72$)) from Dataset 1. p-values and 95% confidence interval were obtained from two-sample t-test. Significant differences ($p < 0.05$) are highlighted with bold text.

| EAA | Reference | Tissue | Sex-adjusted | | Unadjusted | |
|---|---|---|---|---|---|---|
| | | | p | 95% CI | p | 95% CI |
| Horvath AA | normal | healthy | **<0.001** | (2.293, 3.482) | **<0.001** | (2.331, 3.518) |
| | healthy | tumour | **<0.001** | (3.594, 5.972) | **<0.001** | (3.536, 5.919) |
| | healthy | adenoma | 0.631 | (-1.836, 3.009) | 0.656 | (-1.892, 2.987) |
| | normal | tumour | **<0.001** | (6.421, 8.921) | **<0.001** | (6.400, 8.905) |
| | normal | adenoma | **0.006** | (1.022, 5.927) | **0.006** | (1.003, 5.941) |
| | tumour | adenoma | **0.002** | (-6.850, -1.543) | **0.002** | (-6.851, -1.511) |
| Hannum AA | normal | healthy | **<0.001** | (-3.402, -1.532) | **<0.001** | (-3.380, -1.506) |
| | healthy | tumour | **0.046** | (0.020, 2.162) | 0.057 | (-0.031, 2.109) |
| | healthy | adenoma | 0.099 | (-0.308, 3.523) | 0.110 | (-0.364, 3.514) |
| | normal | tumour | **0.042** | (-2.704, -0.048) | **0.038** | (-2.733, -0.075) |
| | normal | adenoma | 0.411 | (-2.925, 1.207) | 0.412 | (-2.957, 1.221) |
| | tumour | adenoma | 0.632 | (-1.613, 2.646) | 0.623 | (-1.614, 2.686) |
| Pheno AA | normal | healthy | **<0.001** | (-11.760, -8.802) | **<0.001** | (-11.686, -8.741) |
| | healthy | tumour | **<0.001** | (-12.629, -8.362) | **<0.001** | (-12.732, -8.482) |
| | healthy | adenoma | 0.695 | (-5.331, 3.573) | 0.666 | (-5.458, 3.506) |
| | normal | tumour | **<0.001** | (-23.238, -18.314) | **<0.001** | (-23.273, -18.368) |
| | normal | adenoma | **<0.001** | (-15.773, -6.546) | **<0.001** | (-15.831, -6.549) |
| | tumour | adenoma | **<0.001** | (4.761, 14.472) | **<0.001** | (4.751, 14.511) |
| SkinBlood AA | normal | healthy | **<0.001** | (-4.263, -2.702) | **<0.001** | (-4.269, -2.708) |
| | healthy | tumour | **<0.001** | (-5.195, -2.459) | **<0.001** | (-5.186, -2.449) |
| | healthy | adenoma | 0.291 | (-3.945, 1.201) | 0.295 | (-3.938, 1.211) |
| | normal | tumour | **<0.001** | (-8.789, -5.830) | **<0.001** | (-8.786, -5.826) |
| | normal | adenoma | **<0.001** | (-7.487, -2.222) | **<0.001** | (-7.486, -2.218) |
| | tumour | adenoma | 0.091 | (-0.401, 5.310) | 0.092 | (-0.403, 5.311) |
| PedBE AA | normal | healthy | 0.093 | (-0.225, 0.017) | 0.095 | (-0.224, 0.018) |
| | healthy | tumour | **<0.001** | (0.161, 0.515) | **<0.001** | (0.160, 0.514) |
| | healthy | adenoma | 0.827 | (-0.487, 0.390) | 0.824 | (-0.487, 0.389) |
| | normal | tumour | **0.019** | (0.039, 0.429) | **0.019** | (0.039, 0.429) |
| | normal | adenoma | 0.499 | (-0.598, 0.294) | 0.499 | (-0.598, 0.294) |
| | tumour | adenoma | 0.101 | (-0.850, 0.077) | 0.102 | (-0.850, 0.078) |
| Wu AA | normal | healthy | **<0.001** | (-1.392, -0.991) | **<0.001** | (-1.387, -0.987) |
| | healthy | tumour | **<0.001** | (-0.771, -0.424) | **<0.001** | (-0.779, -0.431) |
| | healthy | adenoma | 0.296 | (-0.178, 0.577) | 0.316 | (-0.188, 0.574) |
| | normal | tumour | **<0.001** | (-2.041, -1.537) | **<0.001** | (-2.044, -1.540) |
| | normal | adenoma | **<0.001** | (-1.410, -0.574) | **<0.001** | (-1.415, -0.573) |
| | tumour | adenoma | **<0.001** | (0.390, 1.203) | **<0.001** | (0.388, 1.208) |
| Zhang BLUP AA | normal | healthy | **<0.001** | (-5.138, -3.404) | **<0.001** | (-5.123, -3.390) |
| | healthy | tumour | **<0.001** | (-4.380, -2.173) | **<0.001** | (-4.406, -2.200) |
| | healthy | adenoma | 0.349 | (-3.210, 1.149) | 0.341 | (-3.255, 1.141) |
| | normal | tumour | **<0.001** | (-8.857, -6.238) | **<0.001** | (-8.869, -6.251) |
| | normal | adenoma | **<0.001** | (-7.589, -3.014) | **<0.001** | (-7.618, -3.008) |
| | tumour | adenoma | 0.065 | (-0.138, 4.630) | 0.066 | (-0.155, 4.648) |
| Zhang EN AA | normal | healthy | **<0.001** | (-2.057, -0.759) | **<0.001** | (-2.043, -0.746) |
| | healthy | tumour | **0.038** | (0.046, 1.618) | **0.044** | (0.021, 1.595) |
| | healthy | adenoma | 0.597 | (-1.876, 1.086) | 0.575 | (-1.916, 1.070) |
| | normal | tumour | 0.214 | (-1.484, 0.332) | 0.206 | (-1.494, 0.322) |
| | normal | adenoma | **0.023** | (-3.350, -0.256) | **0.023** | (-3.375, -0.258) |
| | tumour | adenoma | 0.133 | (-2.835, 0.380) | 0.135 | (-2.850, 0.389) |
| EpiTOC AA | normal | healthy | **<0.001** | (-0.051, -0.041) | **<0.001** | (-0.051, -0.041) |
| | healthy | tumour | **<0.001** | (-0.069, -0.053) | **<0.001** | (-0.069, -0.053) |
| | healthy | adenoma | 0.254 | (-0.024, 0.006) | 0.256 | (-0.024, 0.006) |
| | normal | tumour | **<0.001** | (-0.116, -0.098) | **<0.001** | (-0.116, -0.098) |
| | normal | adenoma | **<0.001** | (-0.071, -0.039) | **<0.001** | (-0.071, -0.039) |
| | tumour | adenoma | **<0.001** | (0.036, 0.069) | **<0.001** | (0.036, 0.069) |
| HypoClock AA | normal | healthy | **<0.001** | (0.039, 0.047) | **<0.001** | (0.039, 0.047) |
| | healthy | tumour | **<0.001** | (0.052, 0.068) | **<0.001** | (0.052, 0.068) |
| | healthy | adenoma | 0.064 | (-0.001, 0.030) | 0.063 | (-0.001, 0.030) |
| | normal | tumour | **<0.001** | (0.094, 0.112) | **<0.001** | (0.094, 0.112) |
| | normal | adenoma | **<0.001** | (0.042, 0.073) | **<0.001** | (0.042, 0.073) |
| | tumour | adenoma | **<0.001** | (-0.063, -0.028) | **<0.001** | (-0.063, -0.028) |
| MiAge AA | normal | healthy | **<0.001** | (-839.781, -669.545) | **<0.001** | (-840.712, -670.554) |
| | healthy | tumour | **<0.001** | (-1,103.116, -823.647) | **<0.001** | (-1,101.641, -821.904) |
| | healthy | adenoma | 0.342 | (-501.540, 176.483) | 0.346 | (-499.797, 177.613) |
| | normal | tumour | **<0.001** | (-1,876.217, -1559.872) | **<0.001** | (-1,875.719, -1559.093) |
| | normal | adenoma | **<0.001** | (-1,263.955, -570.429) | **<0.001** | (-1,263.199, -570.251) |
| | tumour | adenoma | **<0.001** | (437.366, 1,164.340) | **<0.001** | (437.410, 1163.952) |

**Table S8.** EAA differences between healthy ($n = 715$) and normal ($n = 505$) tissue from Dataset 2. p-values and 95% confidence interval were obtained from two-sample t-test. Significant differences ($p < 0.05$) are highlighted with bold text.

| **EAA** | **Sex-adjusted** | | **Unadjusted** | |
|---|---|---|---|---|
| | **p** | **95% CI** | **p** | **95% CI** |
| Horvath AA | 0.566 | (-0.434, 0.237) | 0.441 | (-0.467, 0.204) |
| Hannum AA | **<0.001** | (0.645, 1.750) | **<0.001** | (0.615, 1.727) |
| Pheno AA | **<0.001** | (0.587, 1.999) | **0.001** | (0.551, 1.983) |
| SkinBlood AA | 0.056 | (-0.014, 1.180) | 0.050 | (-0.001, 1.195) |
| PedBE AA | **0.001** | (0.066, 0.268) | **0.001** | (0.066, 0.267) |
| WuAA | **<0.001** | (0.062, 0.161) | **<0.001** | (0.061, 0.160) |
| Zhang BLUP AA | **0.002** | (0.281, 1.279) | **0.003** | (0.267, 1.267) |
| Zhang EN AA | **<0.001** | (0.603, 1.853) | **<0.001** | (0.577, 1.826) |
| EpiTOC AA | **0.022** | (0.000, 0.004) | **0.020** | (0.000, 0.004) |
| HypoClock AA | **<0.001** | (-0.004, -0.001) | **<0.001** | (-0.004, -0.001) |
| MiAge AA | **0.004** | (8.877, 45.999) | **0.003** | (9.434, 46.576) |

**Table S9.** Coefficients of the fitted linear model (EA C̃A + sex), used in calculating EAA for the first part (for all healthy, normal and tumour samples). Fitted on n=716 healthy samples. For each model term (intercept, age and sex), the table contains coefficient $\beta$, standard error (SE), t-statistic and corresponding p-value.

| **Clock** | **Intercept** | | | | **Age** | | | | **SexM** | | | |
|---|---|---|---|---|---|---|---|---|---|---|---|---|
| | $\beta$ | SE | t | p | $\beta$ | SE | t | p | $\beta$ | SE | t | p |
| Horvath | 23.925 | 1.105 | 21.651 | $< 0.001$ | 0.635 | 0.018 | 34.748 | $< 0.001$ | 1.140 | 0.365 | 3.127 | 0.002 |
| Hannum | 39.919 | 1.908 | 20.918 | $< 0.001$ | 0.439 | 0.032 | 13.907 | $< 0.001$ | 1.648 | 0.630 | 2.616 | 0.009 |
| Pheno | 24.011 | 1.952 | 12.300 | $< 0.001$ | 0.639 | 0.032 | 19.777 | $< 0.001$ | 2.311 | 0.644 | 3.586 | 0.000 |
| SkinBlood | 31.502 | 1.369 | 23.014 | $< 0.001$ | 0.663 | 0.023 | 29.252 | $< 0.001$ | -0.189 | 0.452 | -0.418 | 0.676 |
| PedBE | 6.805 | 0.306 | 22.243 | $< 0.001$ | 0.116 | 0.005 | 22.930 | $< 0.001$ | 0.032 | 0.101 | 0.319 | 0.750 |
| Wu | 8.937 | 0.213 | 41.979 | $< 0.001$ | 0.043 | 0.004 | 12.131 | $< 0.001$ | 0.153 | 0.070 | 2.177 | 0.030 |
| Zhang BLUP | 39.675 | 1.261 | 31.471 | $< 0.001$ | 0.571 | 0.021 | 27.360 | $< 0.001$ | 0.560 | 0.416 | 1.345 | 0.179 |
| Zhang EN | 41.846 | 1.188 | 35.220 | $< 0.001$ | 0.568 | 0.020 | 28.872 | $< 0.001$ | 0.617 | 0.392 | 1.573 | 0.116 |
| EpiTOC | 0.108 | 0.009 | 11.483 | $< 0.001$ | 0.001 | 0.000 | 6.971 | $< 0.001$ | -0.001 | 0.003 | -0.289 | 0.772 |
| HypoScore | 0.872 | 0.004 | 200.53 | $< 0.001$ | 0.000 | 0.000 | -3.108 | 0.002 | -0.003 | 0.001 | -2.007 | 0.045 |
| MiAge | 1,144.41 | 116.12 | 9.855 | $< 0.001$ | 14.43 | 1.922 | 7.510 | $< 0.001$ | -33.26 | 38.32 | -0.868 | 0.386 |

**Table S10.** Coefficients of fitted linear model (EA C̃A + sex), used in calculating EAA for the second part (for all healthy and normal samples). Fitted on n=715 healthy samples. For each model term (intercept, age and sex), the table contains coefficient $\beta$, standard error (SE), t-statistic and corresponding p-value.

| Clock | Intercept | | | | Age | | | | SexM | | | |
|---|---|---|---|---|---|---|---|---|---|---|---|---|
| | $\beta$ | SE | t | p | $\beta$ | SE | t | p | $\beta$ | SE | t | p |
| Horvath | 23.669 | 1.102 | 21.482 | $< 0.001$ | 0.640 | 0.018 | 35.083 | $< 0.001$ | 1.083 | 0.363 | 2.983 | 0.003 |
| Hannum | 39.841 | 1.915 | 20.806 | $< 0.001$ | 0.441 | 0.032 | 13.900 | $< 0.001$ | 1.630 | 0.631 | 2.583 | 0.010 |
| Pheno | 23.443 | 1.939 | 12.090 | $< 0.001$ | 0.649 | 0.032 | 20.209 | $< 0.001$ | 2.183 | 0.639 | 3.416 | 0.001 |
| SkinBlood | 31.127 | 1.361 | 22.868 | $< 0.001$ | 0.669 | 0.023 | 29.692 | $< 0.001$ | -0.273 | 0.449 | -0.608 | 0.543 |
| PedBE | 6.735 | 0.305 | 22.077 | $< 0.001$ | 0.117 | 0.005 | 23.230 | $< 0.001$ | 0.016 | 0.101 | 0.163 | 0.871 |
| Wu | 8.929 | 0.214 | 41.797 | $< 0.001$ | 0.043 | 0.004 | 12.128 | $< 0.001$ | 0.151 | 0.070 | 2.147 | 0.032 |
| Zhang BLUP | 39.392 | 1.257 | 31.326 | $< 0.001$ | 0.576 | 0.021 | 27.656 | $< 0.001$ | 0.496 | 0.414 | 1.197 | 0.232 |
| Zhang EN | 41.642 | 1.188 | 35.048 | $< 0.001$ | 0.571 | 0.020 | 29.044 | $< 0.001$ | 0.571 | 0.392 | 1.459 | 0.145 |
| EpiTOC | 0.104 | 0.009 | 11.262 | $< 0.001$ | 0.001 | 0.000 | 7.450 | $< 0.001$ | -0.002 | 0.003 | -0.551 | 0.582 |
| HypoScore | 0.872 | 0.004 | 200.40 | $< 0.001$ | 0.000 | 0.000 | -3.243 | 0.001 | -0.003 | 0.001 | -1.914 | 0.056 |
| MiAge | 1,107.11 | 115.07 | 9.621 | $< 0.001$ | 15.06 | 1.905 | 7.908 | $< 0.001$ | -41.65 | 37.92 | -1.098 | 0.272 |

**Table S11.** Coefficients of fitted linear model (EA C̃A) and scaling parameters, used in calculating EAAs for the classifier. Linear model was fitted on $n = 341$ healthy samples. Standard Normal distribution scaling parameters were calculated on $n = 556$ samples.

| Clock | Intercept | | | | Age | | | | Scaling parameters | |
|---|---|---|---|---|---|---|---|---|---|---|
| | $\beta$ | SE | t | p | $\beta$ | SE | t | p | mean | SD |
| Horvath | 23.247 | 1.571 | 14.797 | $< 0.001$ | 0.674 | 0.025 | 26.705 | $< 0.001$ | -0.067 | 4.651 |
| Hannum | 42.030 | 2.586 | 16.255 | $< 0.001$ | 0.391 | 0.042 | 9.421 | $< 0.001$ | 1.550 | 9.462 |
| Pheno | 23.452 | 2.845 | 8.243 | $< 0.001$ | 0.653 | 0.046 | 14.293 | $< 0.001$ | 3.199 | 10.792 |
| SkinBlood | 26.171 | 1.712 | 15.285 | $< 0.001$ | 0.733 | 0.028 | 26.638 | $< 0.001$ | -0.523 | 5.423 |
| PedBE | 5.661 | 0.380 | 14.890 | $< 0.001$ | 0.137 | 0.006 | 22.432 | $< 0.001$ | -0.340 | 1.298 |
| Wu | 9.140 | 0.312 | 29.268 | $< 0.001$ | 0.043 | 0.005 | 8.500 | $< 0.001$ | -0.852 | 1.966 |
| Zhang BLUP | 36.325 | 1.524 | 23.834 | $< 0.001$ | 0.617 | 0.024 | 25.191 | $< 0.001$ | -0.827 | 5.692 |
| Zhang EN | 42.729 | 1.620 | 26.374 | $< 0.001$ | 0.569 | 0.026 | 21.846 | $< 0.001$ | -0.765 | 5.383 |
| EpiTOC | 0.049 | 0.010 | 4.817 | $< 0.001$ | 0.002 | 0.000 | 10.871 | $< 0.001$ | -0.007 | 0.032 |
| HypoScore | 0.873 | 0.005 | 180.335 | $< 0.001$ | 0.000 | 0.000 | -2.343 | 0.020 | 0.003 | 0.020 |
| MiAge | 477.101 | 122.258 | 3.902 | $< 0.001$ | 21.746 | 1.964 | 11.070 | $< 0.001$ | -120.004 | 440.154 |

**Table S12.** Folds for cross-validation. Folds in bold were also used for platform-dependent classifier

| Fold | Train | Test |
|---|---|---|
| 1 | GSE101764, GSE149282, GSE142257, GSE166212 | GSE132804_450k, GSE132804_epic |
| 2 | GSE101764, GSE132804_epic, GSE149282, GSE166212 | GSE132804_450k, GSE142257 |
| 3 | GSE101764, GSE132804_epic, GSE142257, GSE166212 | GSE132804_450k, GSE149282 |
| 4 | GSE101764, GSE132804_epic, GSE149282, GSE142257 | GSE132804_450k, GSE166212 |
| 5 | GSE132804_epic, GSE149282, GSE142257, GSE166212 | GSE132804_450k, GSE101764 |
| 6 | GSE101764, GSE149282, GSE132804_450k, GSE166212 | GSE132804_epic, GSE142257 |
| **7** | **GSE101764, GSE132804_450k, GSE142257, GSE166212** | **GSE132804_epic, GSE149282** |
| **8** | **GSE101764, GSE149282, GSE132804_450k, GSE142257** | **GSE132804_epic, GSE166212** |
| **9** | **GSE149282, GSE132804_450k, GSE142257, GSE166212** | **GSE132804_epic, GSE101764** |
| **10** | **GSE132804_epic, GSE149282, GSE132804_450k, GSE166212** | **GSE101764, GSE142257** |
| **11** | **GSE101764, GSE132804_epic GSE132804_450k, GSE166212** | **GSE142257, GSE149282** |
| **12** | **GSE101764, GSE132804_epic, GSE149282, GSE132804_450k** | **GSE142257, GSE166212** |

**Table S13.** Models Coefficients and Performance

| **Model** | Main classifier: $\alpha = 0.05, \lambda = 0.16$ | Parameters: $\alpha = 0.25, \lambda = 0.25$ | Parameters: $\alpha = 0.1, \lambda = 0.35$ | Platform ID included: $\alpha = 0.05, \lambda = 0.68$ |
|---|---|---|---|---|
| **Coefficients** | | | | |
| (Intercept) | -0.7387 | -0.5943 | -0.7094 | -0.5677 |
| Horvath | - | - | - | - |
| Hannum | 0.1601 | 0.0017 | - | 0.0641 |
| Pheno | 0.5132 | 0.3044 | 0.5983 | 0.1729 |
| SkinBlood | -0.0408 | - | - | - |
| PedBE | -0.1533 | -0.0888 | -0.1191 | -0.0881 |
| Wu | -0.5147 | -0.4272 | -0.7161 | -0.2179 |
| Zhang BLUP | -0.0416 | - | - | -0.0190 |
| Zhang EN | -0.1423 | - | -0.0814 | -0.0322 |
| EpiTOC | -0.0351 | - | - | -0.0426 |
| HypoScore | 0.1679 | 0.0212 | 0.1256 | 0.0438 |
| MiAge | -0.1342 | -0.0756 | -0.0602 | -0.0865 |
| sex | 0.4649 | 0.2288 | 0.4171 | 0.1944 |
| **Performance** | | | | |
| ROC-AUC | 0.8858 | 0.8817 | 0.8351 | 0.9207 |
| ROC-AUC 95%CI | [0.8497, 0.9218] | [0.8449, 0.9185] | [0.7910, 0.8793] | [0.8922, 0.9492] |

#### 1.2 Supplementary Figures

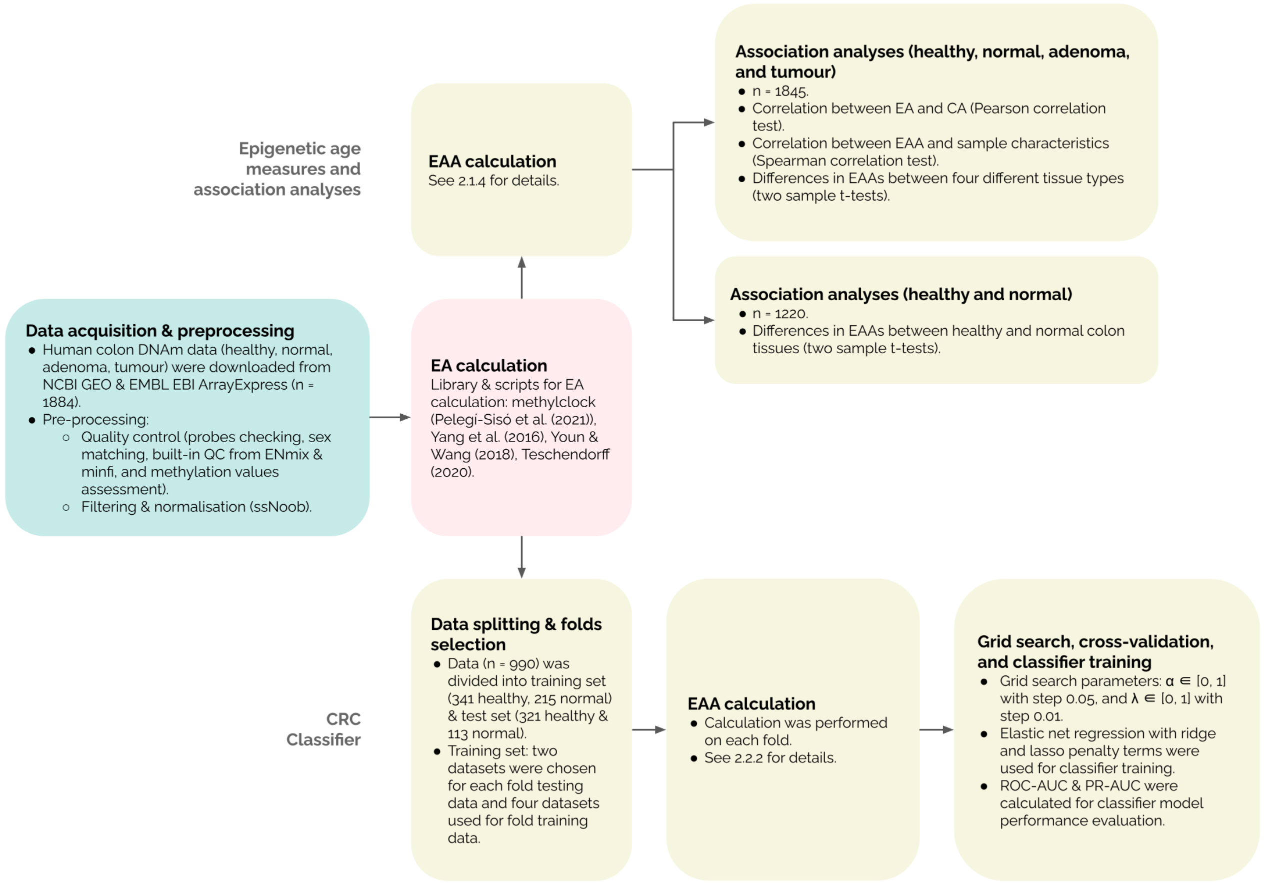


**Figure S1.** Graphical summary of methodology. Abbreviations: CA - Chronological age, DNAm - DNA methylation, EA - Epigenetic age, EAA - Epigenetic age acceleration, F - Female, M - Male, QC - Quality control.

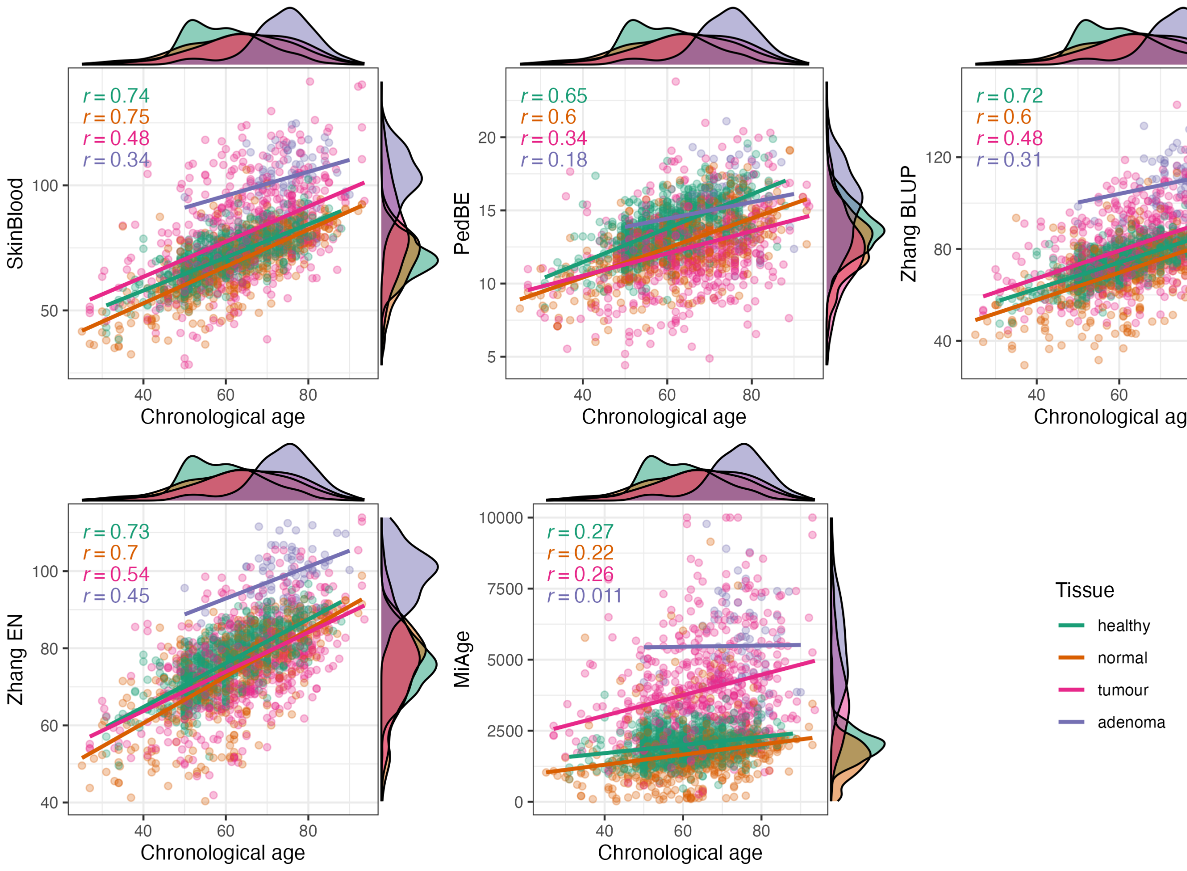


**Figure S2.** Correlation between chronological age and epigenetic age estimates in four different tissues (healthy (n=716), normal (n=522), tumour (n=535), and adenoma (n=72)) based on Pearson correlation test. Different colours represent different tissues.

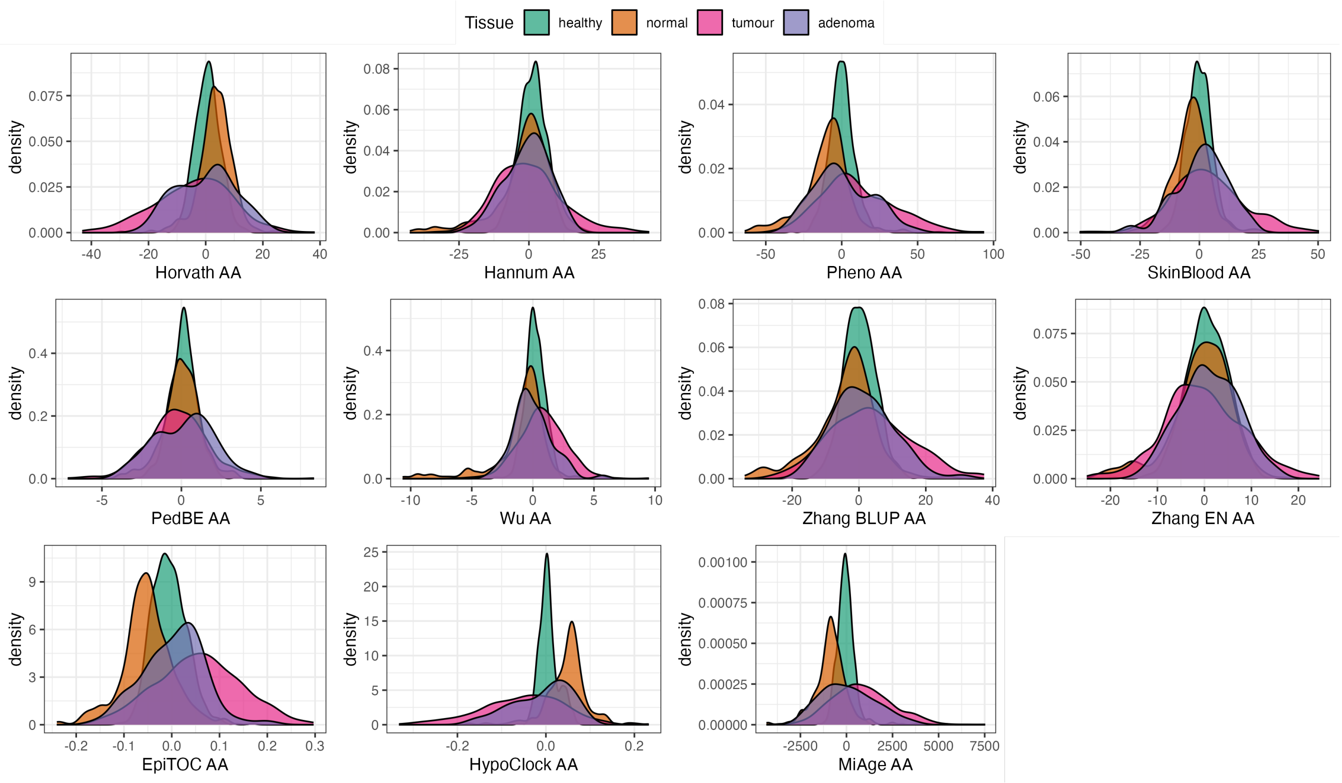


**Figure S3.** Density plots of EAAs distribution in four different tissues (healthy, normal, tumour, adenoma) from Dataset 1.

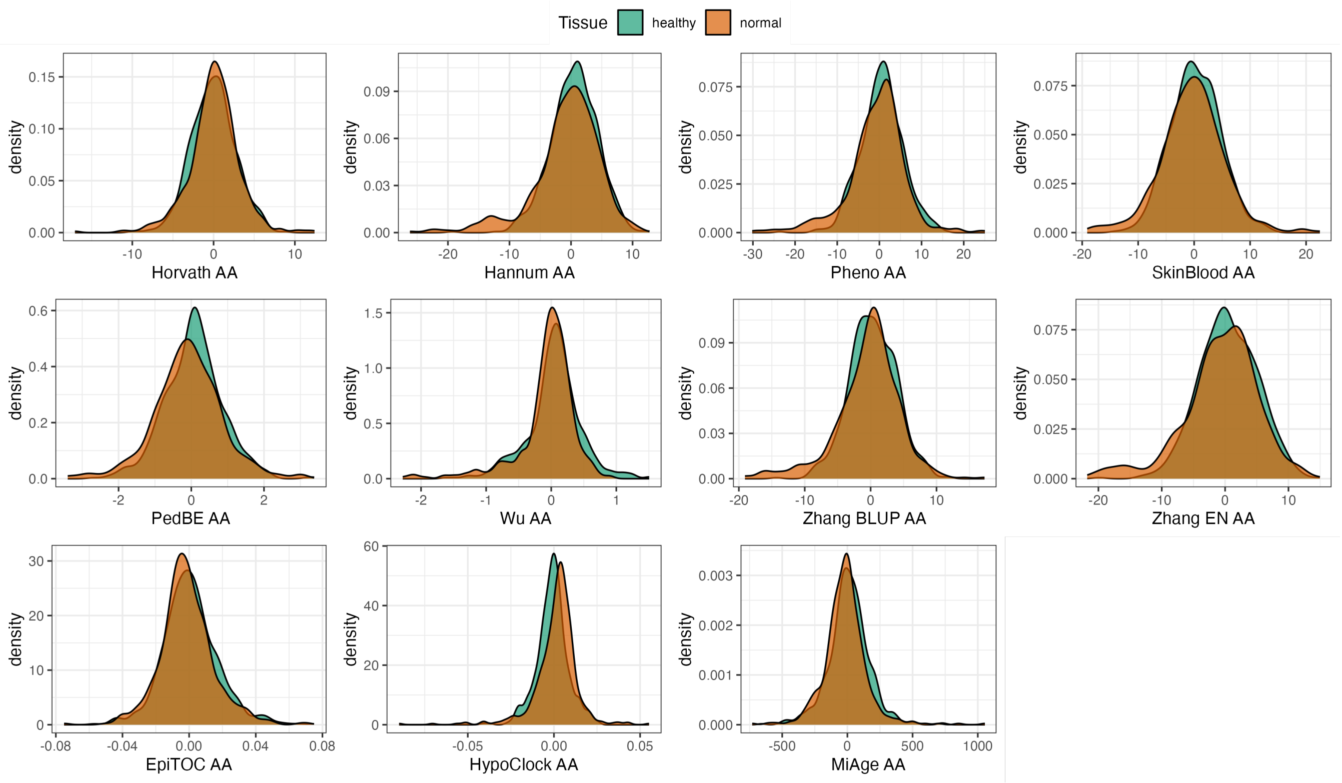


**Figure S4.** Density plots of EAAs distribution in healthy and normal tissues from Dataset 2.

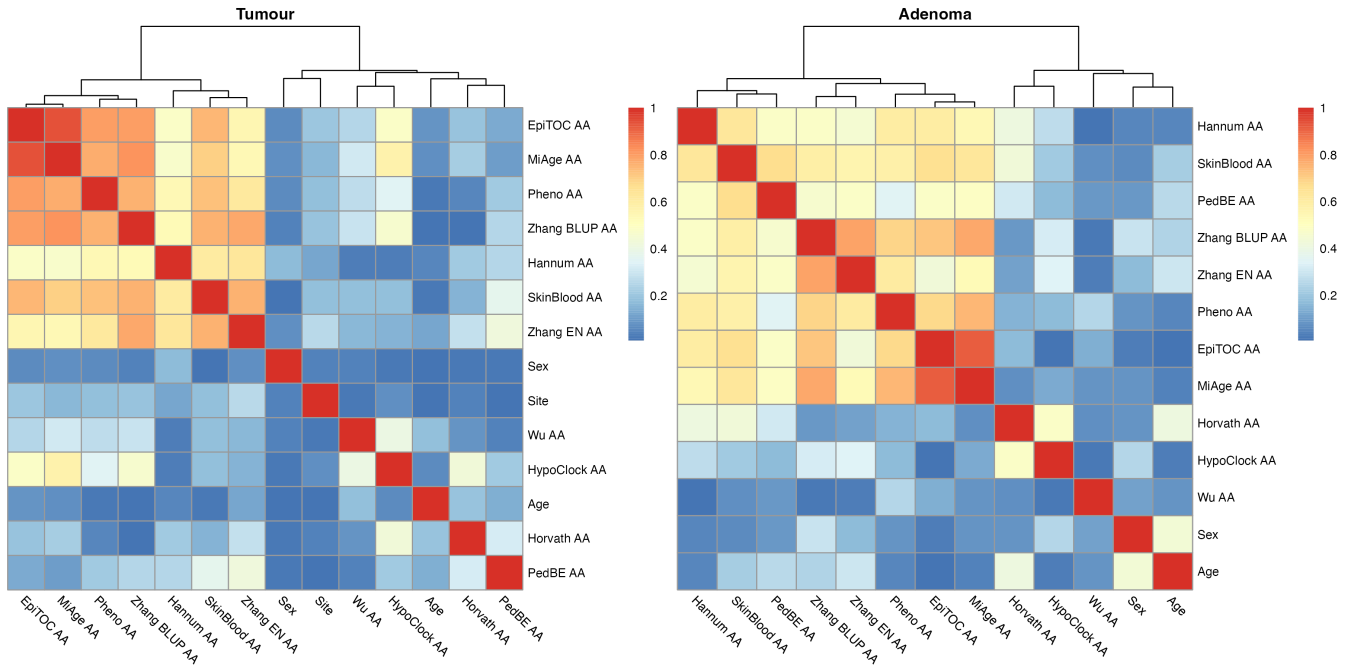


**Figure S5.** Heatmap of Spearman correlation between sample characteristics and sex-adjusted epigenetic age accelerations (EAAs) in tumour and adenoma tissues from Dataset 1. Correlation coefficients are in absolute values.

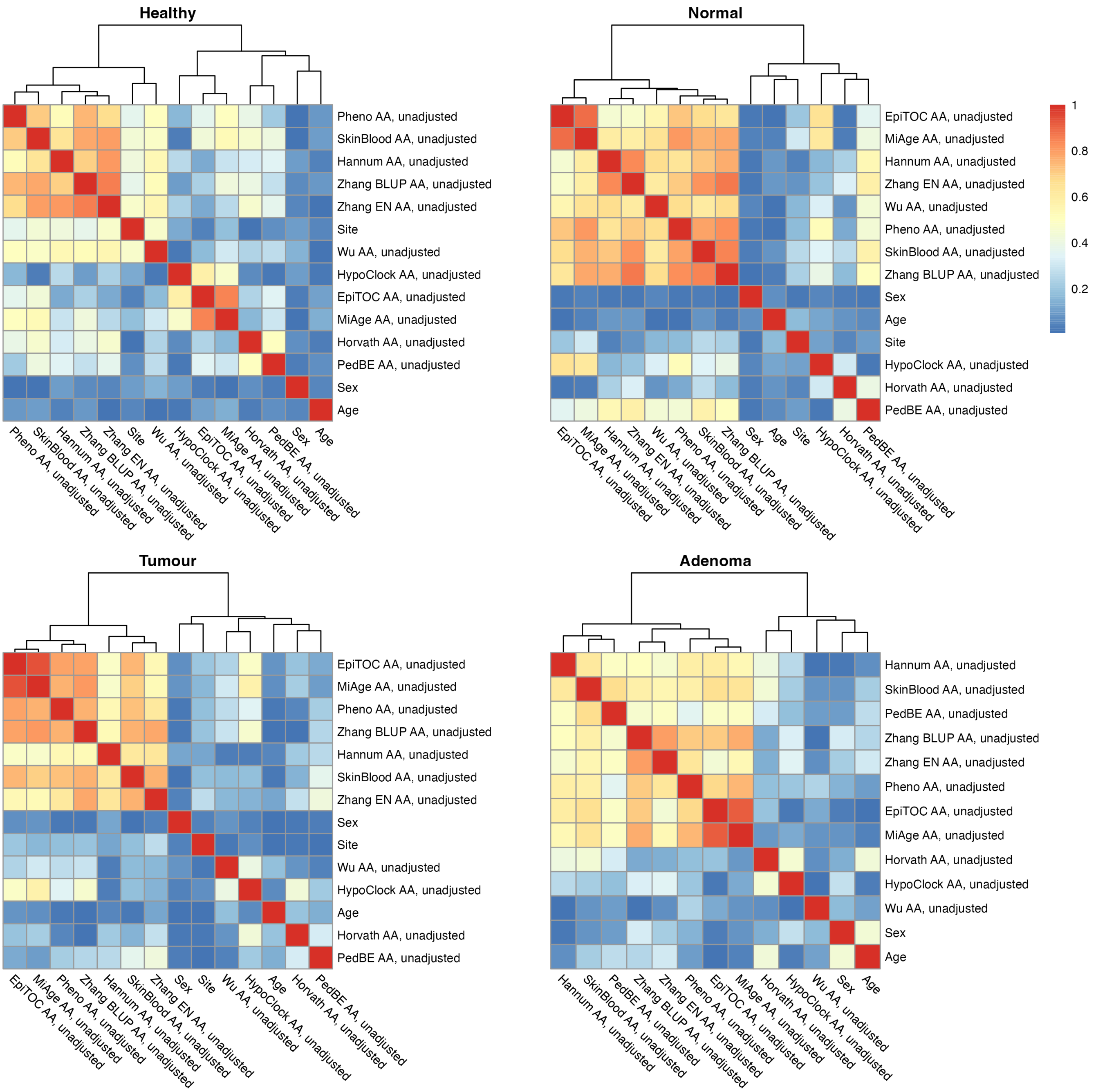


**Figure S6.** Heatmap of Spearman correlation between sample characteristics and unadjusted epigenetic age accelerations (EAAs) in four different tissues (healthy, normal, tumour, and adenoma) from Dataset 1. Correlation coefficients are in absolute values.

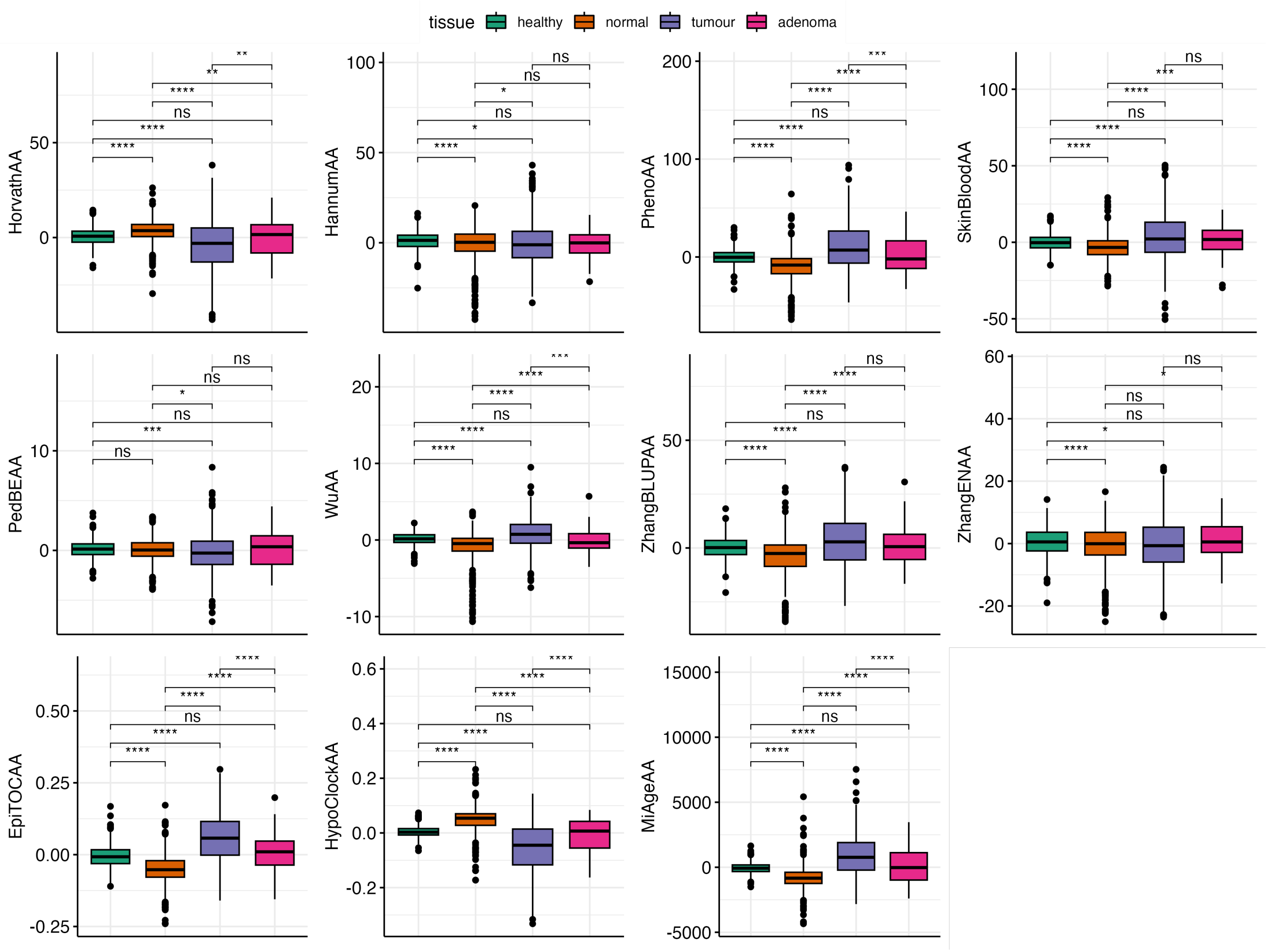


**Figure S7.** Boxplot of sex-adjusted EAAs in four different tissues from Dataset 1. The p-values were obtained from Welch's two-sample t-test. *p<0.05, ** p<0.001, ***p<0.001, ****p<0.0001.

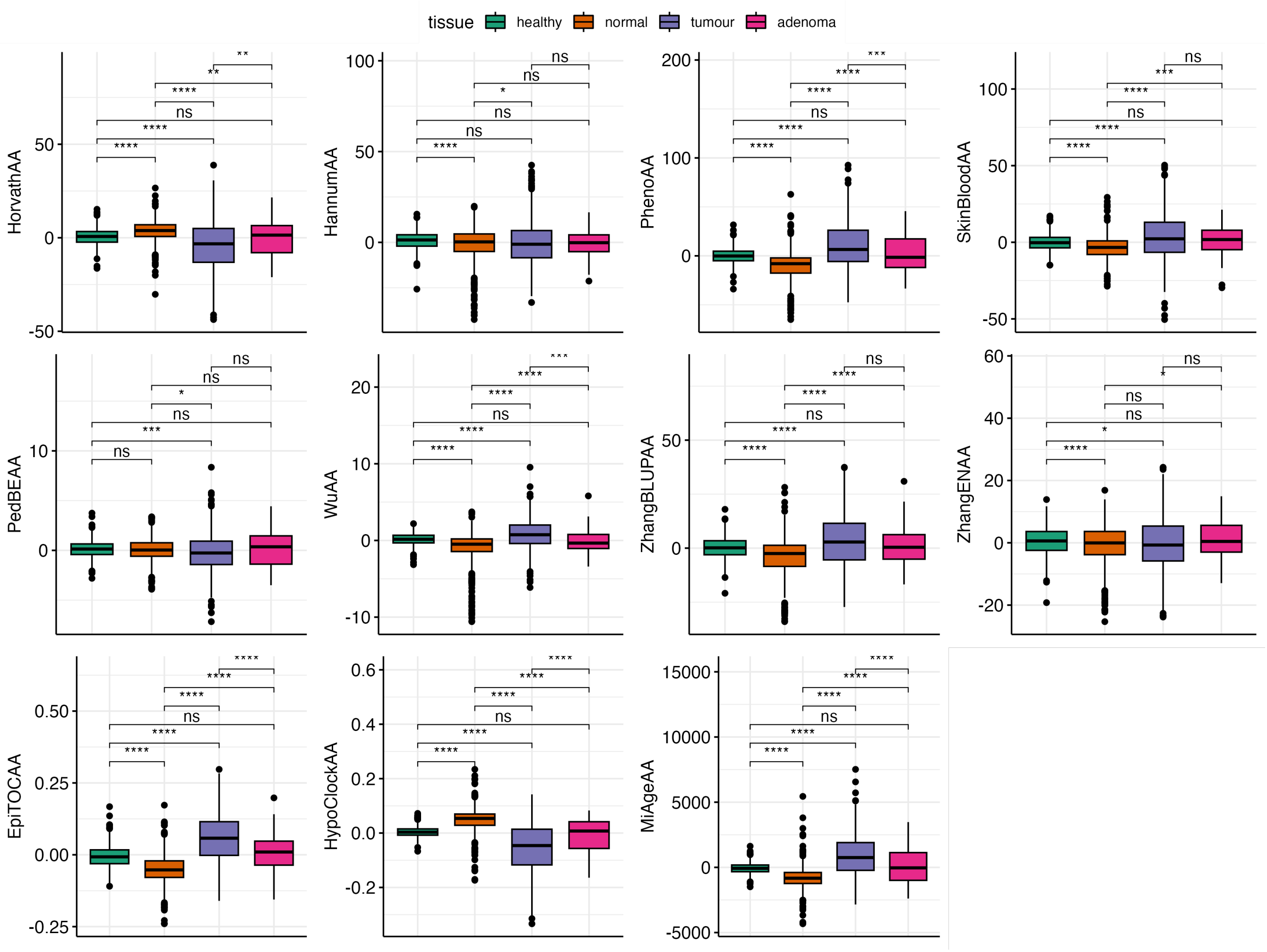


**Figure S8.** Boxplot of unadjusted EAAs in four different tissues from Dataset 1. The p-values were obtained from Welch's two-sample t-test. *p<0.05, ** p<0.001, ***p<0.001, ****p<0.0001.

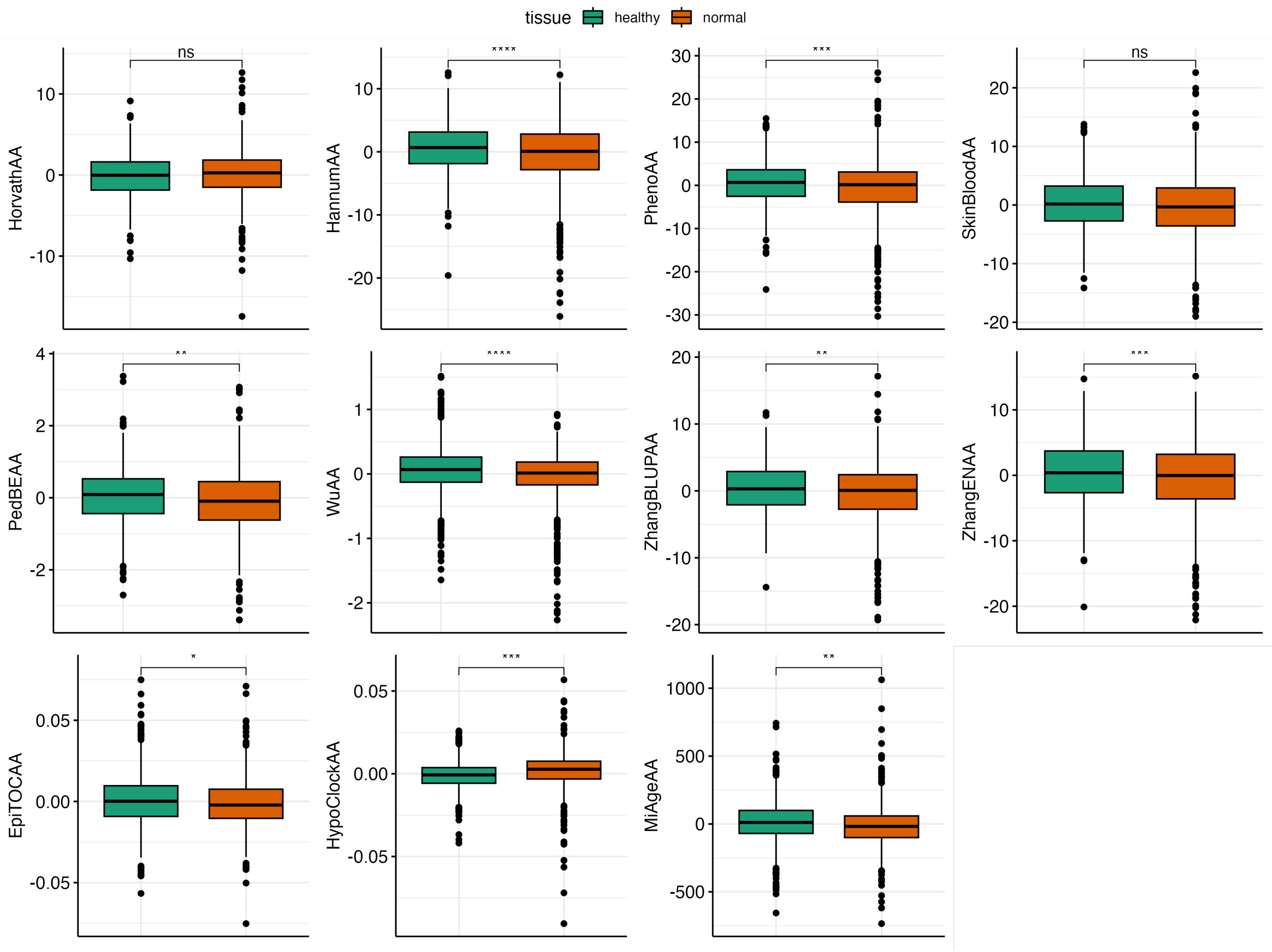


**Figure S9.** Boxplot of unadjusted EAAs in healthy and normal tissues from Dataset 2. The p-values were obtained from Welch's two-sample t-test. *p<0.05, ** p<0.001, ***p<0.001, ****p<0.0001.

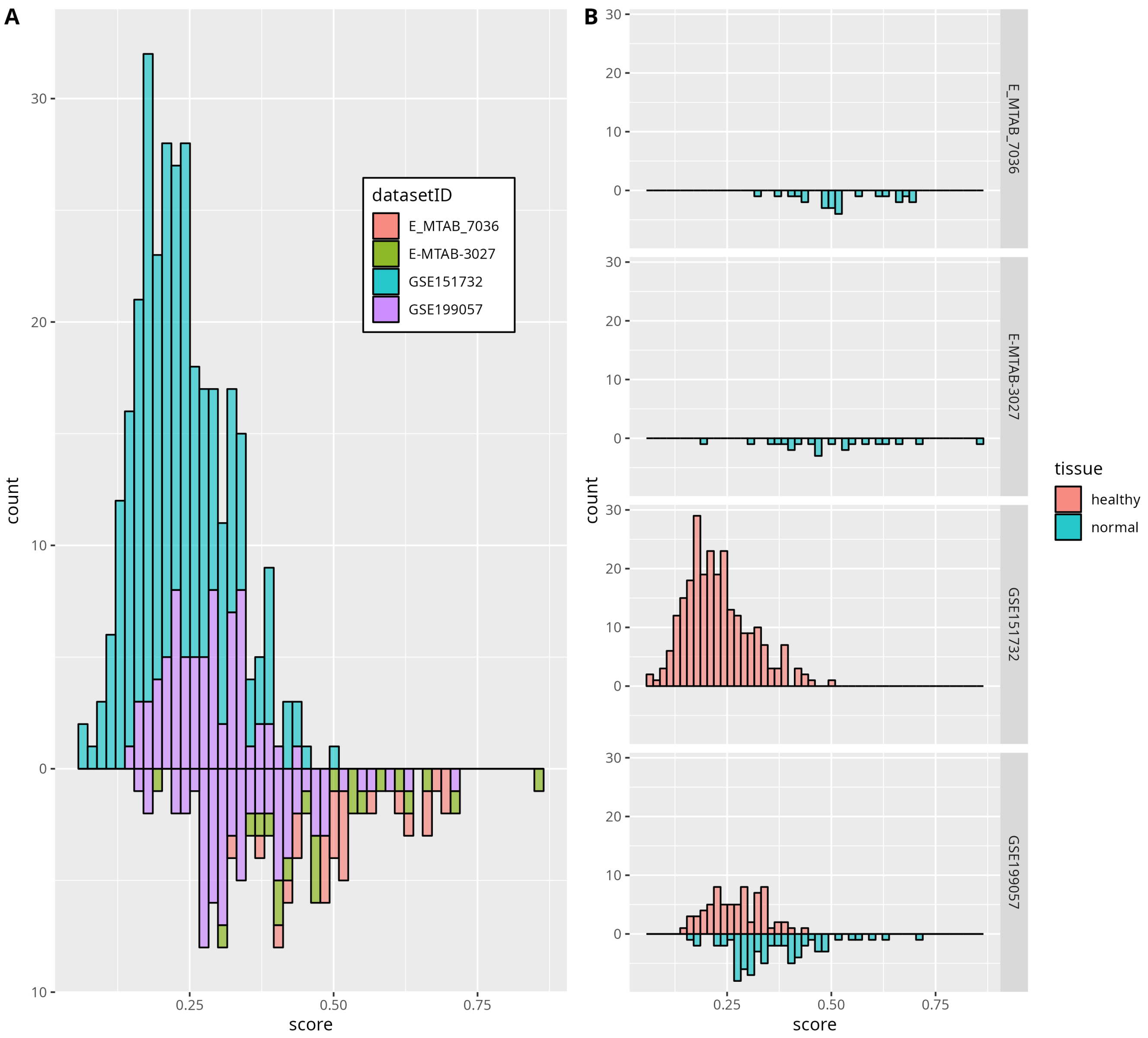


**Figure S10.** Classifier scores histograms for all testing data coloured by dataset (A), and for each dataset separately (B)

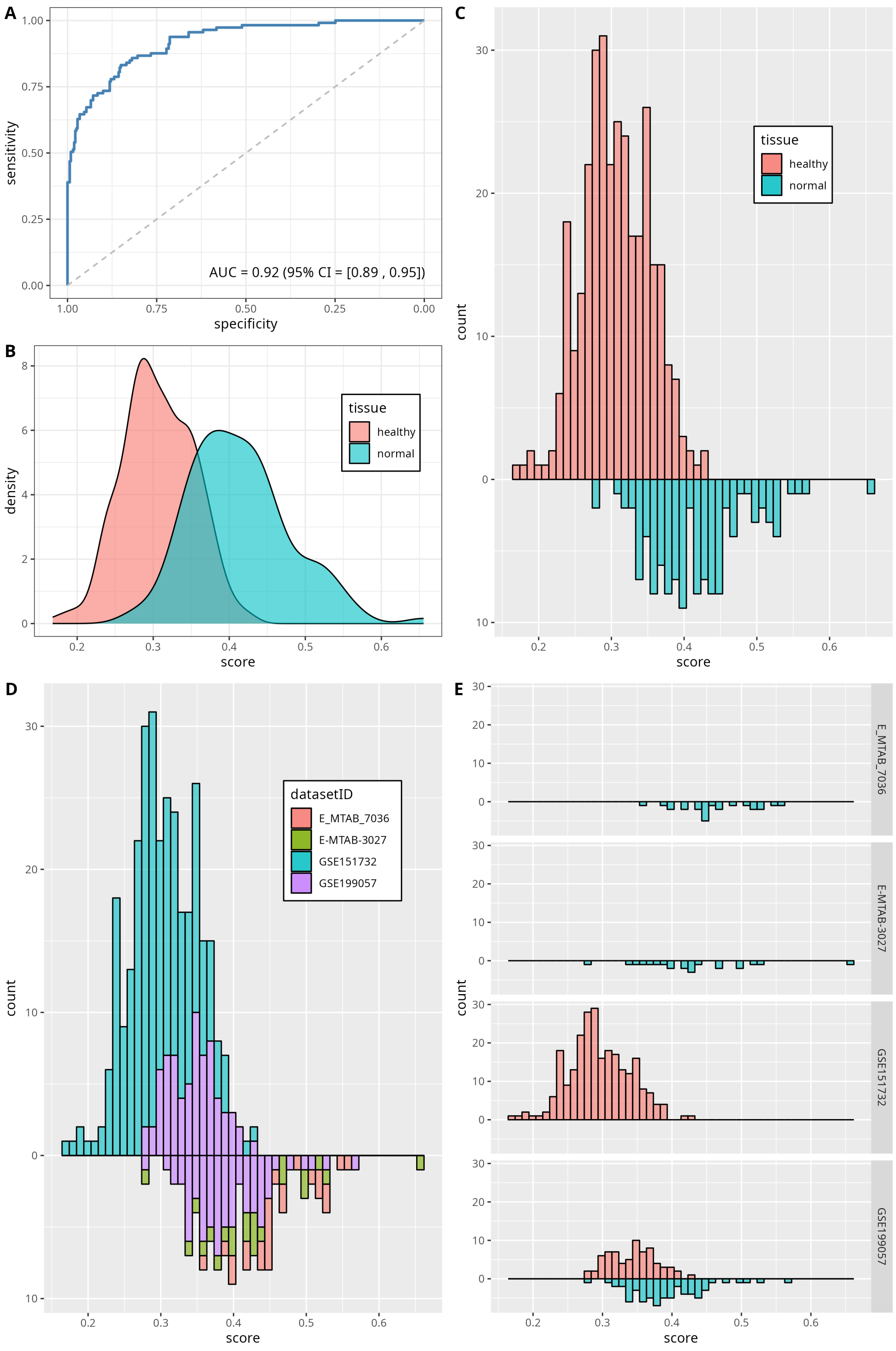


**Figure S11.** Platform-dependent classifier performance. ROC curve (A), density plot (B), scores histograms for all testing data coloured by tissue (C) and dataset ID (D), and for each dataset separately (E)

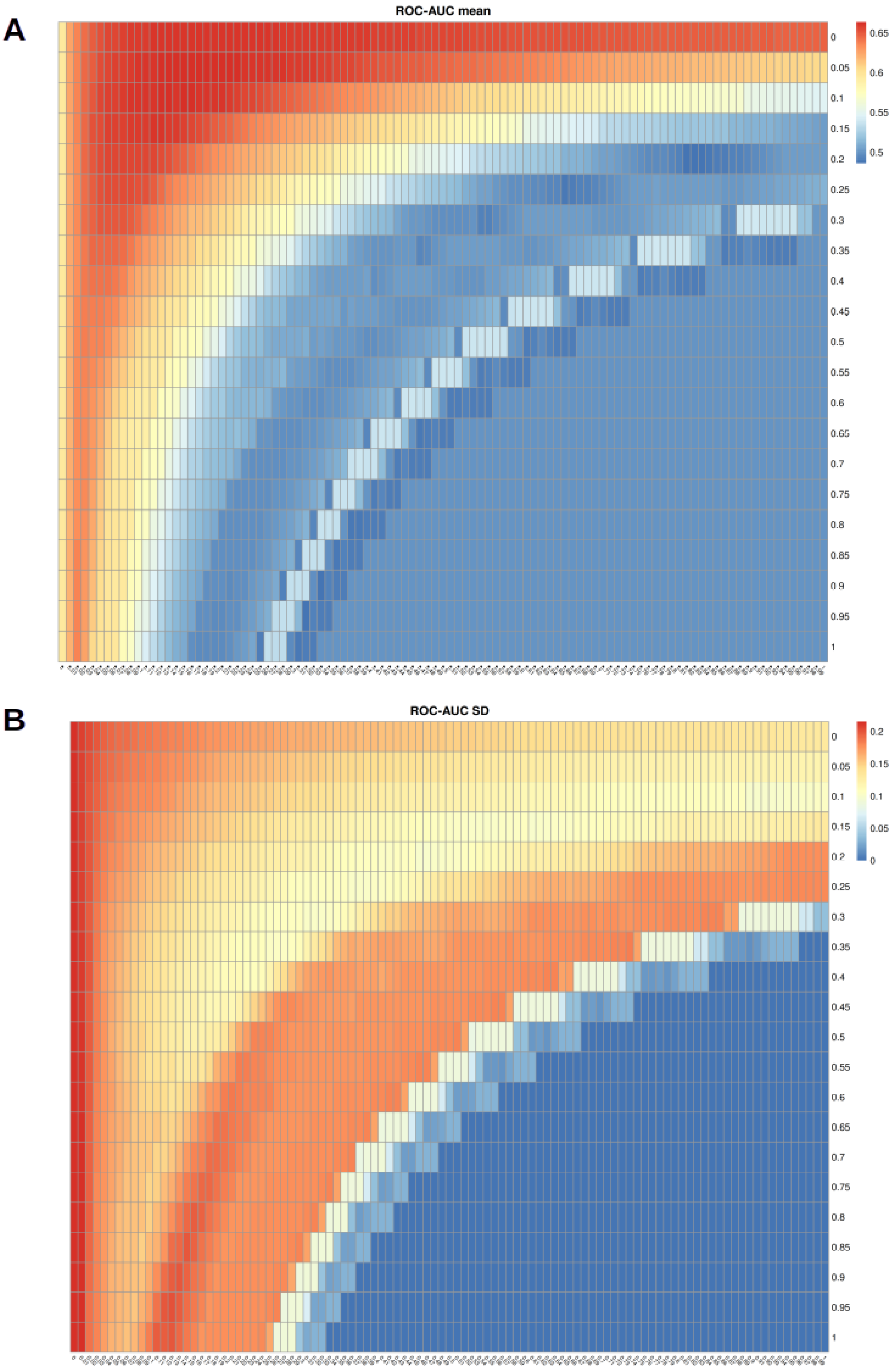


**Figure S12.** Heatmaps for the average ROC-AUC means (A) and standard deviations (B) measures for each pair of parameters $\alpha$ ($y$-axis) and $\lambda$ ($x$-axis) across twelve cross-validation folds.
